# Contrasting patterns of ERK activation in the tail of the striatum in response to aversive and rewarding signals

**DOI:** 10.1101/533299

**Authors:** Giuseppe Gangarossa, Laia Castell, Liliana Castro, Pauline Tarot, Frederic Veyrunes, Pierre Vincent, Federica Bertaso, Emmanuel Valjent

## Abstract

The caudal part of the striatum, also named the tail of the striatum (TS), defines a fourth striatal domain. Determining whether rewarding, aversive and salient stimuli regulate the activity of striatal spiny projections neurons (SPNs) of the TS is therefore of paramount importance to understand its functions, which remain largely elusive. Taking advantage of genetically encoded biosensors (A-kinase activity reporter 3, AKAR3) to record PKA signals and by analyzing the distribution of dopamine D1R- and D2R-SPNs in the TS, we characterized three subterritories: a D2R/A2aR-lacking, a D1R/D2R-intermingled and a D1R/D2R-SPNs-enriched area (corresponding to the amygdalostriatal transition). In addition, we provide evidence that the distribution of D1R- and D2R-SPNs in the TS is evolutionarily conserved (mouse, rat, gerbil). The *in vivo* analysis of extracellular signal–regulated kinase (ERK) phosphorylation in these TS subterritories in response to distinct appetitive, aversive and pharmacological stimuli revealed that SPNs of the TS are not recruited by stimuli triggering innate aversive responses, fasting, satiety or palatable signals whereas a reduction in ERK phosphorylation occurred following learned avoidance. In contrast, D1R-SPNs of the intermingled and D2R/A2aR-lacking areas were strongly activated by both D1R agonists and psychostimulant drugs (d-amphetamine, cocaine, MDMA or methylphenidate), but not by hallucinogens. Finally, a similar pattern of ERK activation was observed by blocking selectively dopamine reuptake. Together, our results reveal that the caudal TS might participate in the processing of specific reward signals and discrete aversive stimuli.

**Abbreviations:** serotonin (5-HT), adenosine A2a receptor (A2aR), A-kinase activity reporter 3 (AKAR3), amygdalostriatal transition area (AST), cyclic adenosine monophosphate (cAMP), dopamine D1 receptor (D1R), dopamine D2 receptor (D2R), dopamine (DA), dopamine- and cAMP-regulated phosphoprotein Mr 32 kDa (DARPP-32), dopamine transporter (DAT), 2,5-dimethoxy-4-iodoamphetamine (DOI), dorsal striatum (DS), enhanced green fluorescence protein (eGFP), extracellular signal-regulated kinase (ERK), high-fat high sugar (HFHS), lateral geniculate nucleus of the thalamus (LGN), medial geniculate nucleus of the thalamus (MGN), 3,4-methylenedioxy methamphetamine (MDMA), norepinephrine (NE), norepinephrine transporter (NET), phencyclidine (PCP), phosphodiesterase type 4 (PDE4), phosphodiesterase type 10 (PDE10A), protein kinase A (PKA), pentylenetetrazole (PTZ), rostral part of the dorsal striatum (rDS), red fluorescent protein (RFP), Research Resource Identifier (RRID), serotonin transporter (SERT), striatal projection neurons (SPNs), substantia nigra pars compacta (SNc), 2,3,5-Trimethyl-3-thiazoline (TMT), tail of the striatum (TS), vesicular glutamate transporter type 2 (VGLUT2), ventral posterior nucleus (VPN).

## Introduction

The striatum, the main input structure of the basal ganglia circuit, controls distinct functions ranging from motor actions to reward processing and motivated behaviors (Nicola 2007; Gerfen and Surmeier 2011; Graybiel and Grafton 2015; Balleine *et al.* 2007). This functional diversity in part relies on its ability to dynamically integrate multimodal excitatory inputs arising from the cortex and the thalamus (Huerta-Ocampo *et al.* 2014). These excitatory inputs are topographically segregated, delineating functional boundaries between the three classical striatal domains that include sensorimotor, associative and limbic domains (Belin *et al.* 2009; Yin and Knowlton 2006; Gruber and McDonald 2012).

However, recent large-scale mouse projectome studies support the existence of a fourth domain (Hunnicutt *et al.* 2016; Menegas *et al.* 2015). Corresponding to the tail of the striatum (TS), this domain is densely innervated by excitatory inputs from the entorhinal, auditory and visual cortices and sensory thalamic nuclei including the lateral and medial geniculate thalamus and the ventral posterior nucleus (Menegas *et al.* 2015; Jiang and Kim 2018; Gangarossa *et al.* 2013b; Hunnicutt *et al.* 2016). The integration of the information arising from these cortical and thalamic areas is modulated by dopaminergic (DA) neurons clustered in the lateral part of the substantia nigra pars compacta (SNc) (Gangarossa *et al.* 2013b; Menegas *et al.* 2015; Poulin *et al.* 2018). These DA neurons, which express VGLUT2 and lack D2 autoreceptors (Gangarossa *et al.* 2013b; Poulin *et al.* 2018), contribute to reinforcement learning that promotes avoidance of threatening stimuli (Menegas *et al.* 2018).

In addition to its specialized inputs connectivity, the TS is also characterized by a topographic non-random distribution of striatal projection neurons (SPNs) expressing D1R (D1R-SPNs) and D2R/A2aR (D2R/A2aR-SPNs). Indeed, we have previously identified in the TS a territory lacking D2R/A2aR-SPNs and exclusively composed of D1R-SPNs (Gangarossa *et al.* 2013b). Although this peculiar organization suggests that information processed within the TS could differ from the one described in the sensorimotor, associative and limbic domains, the nature of the stimuli that recruit the SPNs of the TS remain largely unknown.

The present study aims to determine whether stimuli of different nature (rewarding, aversive and salient) modulate the activation of SPNs in the TS. We first performed an accurate characterization of the distribution of D1R- and D2R/A2aR-SPNs within the TS, which allowed us to delineate three striatal subterritories, namely the (*i*) intermingled area, the (*ii*) D2R/A2aR-lacking area and the (*iii*) amygdalostriatal transition (AST) area. We also analyzed the ability of distinct appetitive, aversive and pharmacological stimuli to activate caudal SPNs by monitoring ERK phosphorylation (P-ERK), used here as a molecular readout of striatal activation (Bertran-Gonzalez *et al.* 2010; Hutton *et al.* 2017; Gangarossa *et al.* 2013c; Valjent *et al.* 2019). Our results reveal opposing patterns of ERK activation in the TS in response to innate or learned avoidance behaviors compared to rewarding experiences evoked by psychostimulant drugs.

## Materials and methods

### Animals

C57Bl/6J (n = 153) (RRID:IMSR_JAX:000664 from Charles River Laboratories, France, RRID:SCR_003792), FVB/N (n = 3) (RRID:IMSR_JAX:001800 from Janvier Laboratories, France), *Drd2-*eGFP (n = 53) (RRID:MMRRC_000230-UNC), *Nr4a1*-eGFP (n = 7) (RRID:MMRRC_012015-UCD) (Gong *et al.* 2003) and *Drd2-*eGFP/*Drd1a-*tdTomato (n = 4, kindly provided by C. Kellendonk, Columbia University, NY USA and previously characterized (Biezonski *et al.* 2015)) mice were used in the present study. *Drd2-*eGFP/*Drd1a-*tdTomato mice were generated by crossing *Drd1a*-tdTomato (Tg(*Drd1a*-tdTomato)6Calak/J, RRID:IMSR_JAX:016204) and *Drd2*-eGFP (Tg(*Drd2*-EGFP)S118Gsat/Mmnc, RRID:IMSR_MMRRC:000230) mice. Only male 8-10 week-old mice (25-30 grams) were used in the current study. Unless specifically stated, all animals (mice, rats, gerbils) were housed in groups of 2 to 5 per cage (standard sizes according to the European animal welfare guidelines 2010/63/EC) and maintained in a 12 h light/dark cycle (lights on from 7:30 am to 7:30 pm), in stable conditions of temperature (22°C) and humidity (60%), with food and water provided *ad libitum*. This study was not pre-registered.

We used PCR to determine the genotype of our transgenic mice. The following PCR primers were used: 5’-GCAGAAGAACGGCATCAAG-3’ (forward) and 5’-GCTCAGGTAGTGGTTGTCG-3’ (reverse) for GFP; 5’-CTTCTGAGGCGGAAAGAAC-3’ (forward) and 5’-TTTCTGATTGAGAGCATTCG-3’ (reverse) for tdTomato. 18 adult males from nine different wild mouse taxa (genus *Mus*), which included 3 subspecies of *Mus musculus* [*Mus musculus domesticus* (n = 2), *Mus musculus castaneus* (n = 2) and *Mus musculus musculus* (n = 2)], 4 species of the subgenus *Mus* [*Mus spretus* (n = 2), *Mus spicilegus* (n = 2), *Mus macedonicus* (n = 2) and *Mus caroli* (n = 2)] and 2 species from other *subgenera* (*Nannomys minutoides* (n = 2) and *Coelomys pahari* (n = 2)), were purchased from the Wild Mouse Genetic Repository of Montpellier (France) except for *Nannomys minutoides* that were raised at the breeding facility (CECEMA) of the University of Montpellier, France. Male 3-4 month-old Mongolian gerbils (n = 3, 70-80 grams) (Janvier Laboratories, France) were kindly provided by Gilles Desmadryl (INM Montpellier France). Male 10 week-old Sprague-Dawley rats (n = 3, 450 grams, RRID:RGD_734476) were purchased from Charles River Laboratories (RRID:SCR_003792, France). All experiments were performed during the light phase (notably, 10 am to 4 pm). Male wild mouse taxa, gerbils and rats were used exclusively for the neuroanatomical characterization (Fig. 1). For behavioral and pharmacological experiments arbitrary assignments were used to allocate mice to specific groups. No randomization was performed to allocate subjects in the study. No blinding was performed for animal experiments. Housing and experimental procedures were approved by the French Agriculture and Forestry Ministry (C34-172-13). Experiments were performed in accordance with the animal welfare guidelines 2010/63/EC of the European Communities Council Directive regarding the care and use of animals for experimental procedures. All animals were anaesthetized (euthasol, 360 mg/kg, i.p., TVM lab, France) immediately after acute pharmacological or behavioral experiments (see *Tissue preparation and immunofluorescence* section) (Fig. 1). None of these manipulations required painful and invasive techniques. Male wild mouse taxa, gerbils and rats were immediately anaesthetized (euthasol, 360 mg/kg, i.p.) in their home cages (no pharmacological or behavioral experiments) before transcardiac perfusion with 4% (weight/vol) paraformaldehyde and exclusively used for the neuroanatomical characterization (see Fig. 1, Fig. 4 and *Tissue preparation and immunofluorescence* section). No invasive or painful techniques (surgery) were used during our manipulations.

**Figure 1:**
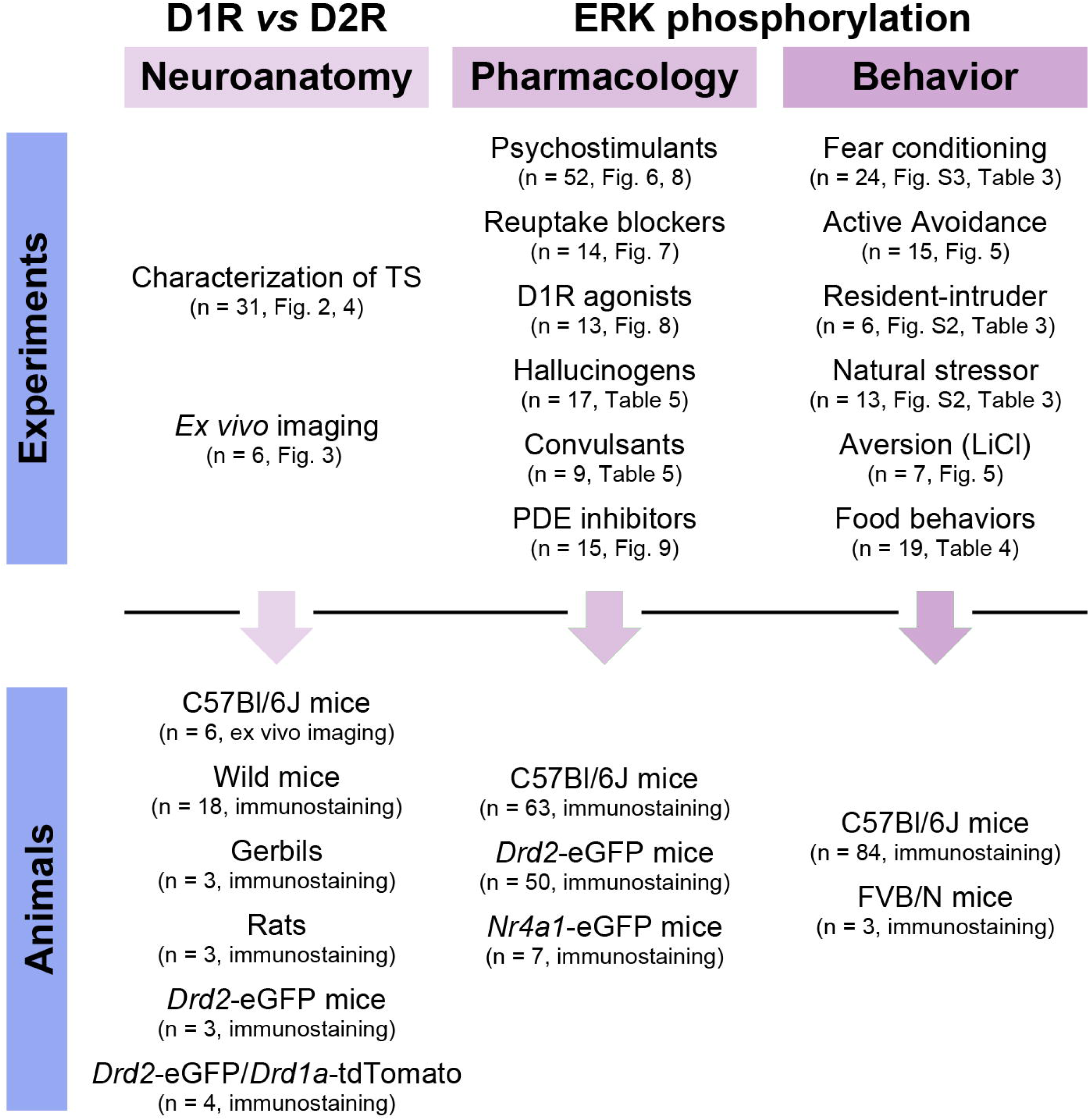
Description of the study. The whole study was designed in order to characterize the cell-type specific organization (Neuroanatomy) as well as the molecular patterns of ERK phosphorylation (Pharmacology and Behavior) within the territories of the tail of the striatum (TS). The figure summarizes the experimental conditions and the animal strains used for each purpose (n = number of mice). For the detailed repartition of the experimental groups please see Figures, Tables and Supplemental Tables. For the detailed description of each experiment please see Material and Methods. No animals were excluded in this study. D1R: dopamine D1 receptor; D2R: dopamine D2 receptor; ERK: extracellular signal–regulated kinase; LiCl: lithium chloride; PDEs: phosphodiesterases; TS: tail of the striatum.

### Drugs and treatment

Drugs and treatment are detailed in Table 1. All drugs were acquired within the last 5 years and stored following the manufacturers recommendation. Drugs were administered at doses in agreement with previous studies showing behavioral and/or molecular modifications (see Table 1). For *in vivo* pharmacological experiments, mice were habituated to handling and saline injections three consecutive days before the experiments. Drugs were dissolved in 0.9% (w/v) NaCl (saline) and administered on day 4. All the mice were injected in the home cage and perfused 15 or 30 min after injection.

**Table 1.** List of drugs.

| Drugs | Doses | Catalog N | Supplier (RRID) | References |
| --- | --- | --- | --- | --- |
| d-amphetamine | 10 mg/kg, i.p. | #2813 | RRID:SCR_003689 | (Gangarossa *et al.* 2013a) |
| Cocaine | 15 mg/kg, i.p. | #C5776 | RRID:SCR_008988 | (Gangarossa *et al.* 2013a) |
| MDMA | 10 mg/kg, i.p. | #M5029 | RRID:SCR_008988 | (Doly *et al.* 2009) |
| Methylphenidate | 15 mg/kg, i.p. | #M2892 | RRID:SCR_008988 | (Pascoli *et al.* 2005) |
| GBR12783 | 10 mg/kg, i.p. | #0513 | RRID:SCR_003689 | (Valjent *et al.* 2010) |
| Desipramine | 20 mg/kg, i.p. | #3067 | RRID:SCR_003689 | (Valjent *et al.* 2004) |
| Fluoxetine | 10 mg/kg, i.p. | #0927 | RRID:SCR_003689 | (Valjent *et al.* 2004) |
| SKF81297 | 5 mg/kg, i.p. | #1447 | RRID:SCR_003689 | (Gangarossa *et al.* 2013c) |
| SKF83822 | 5 mg/kg, i.p. | #2075 | RRID:SCR_003689 | (Gangarossa *et al.* 2011) |
| SKF83959 | 5 mg/kg, i.p. | #2074 | RRID:SCR_003689 | (Gangarossa *et al.* 2011) |
| Apomorphine | 3 mg/kg, s.c. | #2073 | RRID:SCR_003689 | (Gangarossa *et al.* 2013c) |
| SCH23390 | 0.1 mg/kg, i.p. | #0925 | RRID:SCR_003689 | (Valjent *et al.* 2000) |
| LiCl | 150 mg/kg, i.p. | #L4408 | RRID:SCR_008988 | (Longoni *et al.* 2011) |
| DOI | 10 mg/kg, i.p. | #D101 | RRID:SCR_008988 | (Halberstadt *et al.* 2009) |
| Phencyclidine | 3 mg/kg, s.c. | #P3029 | RRID:SCR_008988 | (Mouri *et al.* 2012) |
| Dizocilpine | 0.3 mg/kg, i.p. | #0924 | RRID:SCR_003689 | (Valjent *et al.* 2000) |
| PTZ | 60 mg/kg, i.p. | #P6500 | RRID:SCR_008988 | (Assaf *et al.* 2011) |
| Kainate | 20 mg/kg, i.p. | #K0250 | RRID:SCR_008988 | (Blüthgen *et al.* 2017) |
| Papaverine | 30 mg/kg, i.p. | #P3510 | RRID:SCR_008988 | (Nishi *et al.* 2008) |
| Rolipram | 10 mg/kg, i.p. | #0905 | RRID:SCR_003689 | (Nishi *et al.* 2008) |
| SKF38393 | 1 μM | #0922 | RRID:SCR_003689 | (Polito *et al.* 2015) |
| CGS21680 | 1 μM | #1063 | RRID:SCR_003689 | (Polito *et al.* 2015) |
| Forskolin | 12.5 μM | #1099 | RRID:SCR_003689 | (Polito *et al.* 2015) |

d-amph, d-amphetamine; MDMA, 3,4-Methylenedioxymethamphetamine; LiCl, Lithium chloride; DOI, 2,5-Dimethoxy-4-iodoamphetamine; Dizocilpine, MK801; PTZ, Pentylene-tetrazol

### Animal behaviors

#### Auditory fear conditioning

The experiments were carried out in a fear conditioning apparatus comprising two test boxes placed within ventilated and soundproof cubicles (Imetronic, Pessac, France). Two different contextual configurations were used for the conditioning (box A: smooth white foam on the walls, square cage, metal grid, smooth floor; box B: dark honeycomb foam on the walls, circle cage, metal grid, rough floor). Boxes were cleaned before and after each session with 70% ethanol. After 10 min of habituation to box A or B, mice received 5 pairings of one tone (CS+, 10 s) with an unconditioned stimulus (US: 0.6 mA scrambled footshock, 2 s, coinciding with the last 2 s of CS+ presentation). The interval between each pairing was arbitrarily assigned between 35 and 60 s. The session finished 30 seconds after the end of the last CS+. Freezing behavior was recorded using a tight infrared frame and the threshold for considering freezing behavior was set up at 2 seconds. The first 10 min of habituation were used to assess the basal freezing. Mice were perfused immediately after conditioning.

#### Active avoidance

The experiments were carried out in a shuttle box placed within a ventilated and soundproof cubicle (Imetronic, Pessac, France). The apparatus consisted in two square compartments (20 cm × 20 cm) separated by a partition. Each compartment was equipped with metal grid, independent houselights and tight infrared beams frame to measure locomotor activity. Between each session the apparatus was cleaned with 70% ethanol. Mice were first subjected to a habituation session during which they were exposed to 10 alternating presentations of two different tones (8 s). The two tones were stopped as soon as mice moved to the adjacent compartment. A conditioning session (30 min) comprised 30 training trials. A training trial consisted of an 8 s avoidance interval, followed by a 5 s escape interval. During the avoidance interval, a tone (CS+) was presented for the duration of the interval or until the mice shuttled (avoidance behavior) to the other compartment. If the animal avoided, the CS+ was terminated and no escape interval was presented. If the animal did not avoid, an unconditioned stimulus (US: 0.7 mA scrambled footshock coinciding with the last 5 s of CS+ presentation) was delivered through the grid floor of the occupied half of the shuttle box. In such paradigm, the US was used to motivate the animal to move to the other compartment (escape), thus terminating the trial. During the intertrial interval, the animal awaited the next trial and was free to cross between compartments at will. During the conditioning session mice were also randomly exposed to the other tone (CS-, 8 s), which was never associated with the US. The percentage of avoidances was used as an index of the learning performance. Mice were perfused immediately after the habituation to the shuttle box, the first or the second conditioning session.

#### Resident-intruder

This test is based on territory defensive behaviors against unfamiliar intruding conspecifics. A single-housed resident male (3-to 6-month-old sexually-experienced male FVB/N, 40 g) was confronted in its home-cage to a group-housed intruder male C57Bl/6J mouse for 10 min. Both the resident and intruder were perfused immediately after the test.

#### Natural stressors

The mice were habituated in a medium-sized box during the first day. Twenty-four hours later each animal was transferred to the same cage. After 15 min of habituation, a petri dish containing either mouse’s own bedding (control condition) or rat bedding was introduced in the cage. Similarly, after 15 min of habituation mice were exposed to a filter paper impregnated with water (control condition) or 20 μl of TMT (2,3,5-Trimethyl-3-thiazoline, a constituent of fox urine). Rat bedding and TMT represent two natural stressors (predator odors). All mice were perfused immediately after 15 min of exposure to mouse bedding, rat bedding, water or TMT.

#### Food-related behavior

Mice were single-housed during the whole protocol. Five groups of mice were used: (1) a control group with *ad libitum* access to chow pellets, (2) a fasted group food-deprived for 12 hours (dark period), (3) a group of fasted/refed mice (one cycle), (4) a group of fasted/refed mice during six cycles and (5) a group of bingeing mice daily exposed to high-fat high-sugar (HFHS) diet during 1h/day for six intermittent days. All mice were perfused with PFA 4% 30 min following the last exposure to food (chow or HFHS diet).

### Tissue preparation and immunofluorescence

All animals were rapidly anaesthetized with euthasol (360 mg/kg, i.p., TVM lab, France) and transcardially perfused with 4% (weight/vol) paraformaldehyde in 0.1 M sodium phosphate buffer (pH 7.5). Euthasol (360 mg/kg, i.p), a pentobarbital sodium solution, allows rapid, efficient and deep anesthesia necessary to preserve phosphorylation. For the neuroanatomical characterization mice were perfused with no specific order. For the pharmacological and behavioral experiments mice were perfused every 6 minutes by arbitrarily alternating subjects and groups (behavioral conditions or pharmacological treatments). Brains were post-fixed overnight in the same solution and stored at 4°C. Tissues were prepared as previously described (Gangarossa *et al.* 2013a). Thirty μm-thick sections were cut with a vibratome (Leica, France, RRID:SCR_008960) and stored at −20°C in a solution containing 30% (vol/vol) ethylene glycol, 30% (vol/vol) glycerol, and 0.1 M sodium phosphate buffer, until they were processed for immunofluorescence. Striatal sections were identified using the mouse brain atlas (Franklin, K. and Paxinos, G. 2008) and sections comprised between -1.40 to -1.70 mm from bregma were included in the analysis. Sections were processed as follows: free-floating sections were rinsed three times (10 minutes/wash) in Tris-buffered saline (TBS, 50 mM Tris–HCL, 150 mM NaCl, pH 7.5). After 15 minutes of incubation in 0.2% (vol/vol) Triton X-100 in TBS, sections were rinsed again in TBS and then blocked for 1 hour in a solution of 3% BSA in TBS. Sections were then incubated overnight or for 72 hours at 4°C with the primary antibodies (Table 2) diluted in a TBS solution containing 1% BSA and 0.15% Triton X-100. Sections were rinsed three times for 10 minutes in TBS and incubated for 45 minutes with goat Cy3-coupled anti-rabbit (1:500, Jackson ImmunoResearch Labs Cat# 111-165-003, RRID:AB_2338000), goat Alexa Fluor 647-coupled anti-guinea pig (1:500, Molecular Probes, Cat# A-21450, RRID:AB_141882) and goat Alexa

**Table 2.** List of primary antibodies.

| **Antigen** | **Host** | **Dilution** | **Supplier** | **Catalog N°** | **RRID** |
| --- | --- | --- | --- | --- | --- |
| eGFP | Chicken | 1:1000 | Life Technologies | A10262 | RRID:AB_2534023 |
| RFP | Rabbit | 1:1000 | MBL | PM005 | RRID:AB_591279 |
| DARPP-32 | Rabbit | 1:1000 | Cell Signaling | #2306 | RRID:AB_823479 |
| D2R | Guinea Pig | 1:1000 | Frontier Institute | D2R-GP-Af500 | RRID:AB_2571597 |
| A2aR | Guinea Pig | 1:1000 | Frontier Institute | A2A-GP-Af1000 | RRID:AB_2571656 |
| D1R | Guinea Pig | 1:1000 | Frontier Institute | D1R-GP-Af500 | RRID:AB_2571595 |
| ERK1/2 | Rabbit | 1:500 | Cell Signaling | #4695 | RRID:AB_390779 |
| P-ERK1/2 | Rabbit | 1:500 | Cell Signaling | #9101 | RRID:AB_331646 |

eGFP, enhanced green fluorescent protein; RFP, red fluorescent protein; DARPP-32, Dopamine- and cAMP-regulated phosphoprotein, Mr 32 kDa; D2R, dopamine D2 receptor; A2aR, adenosine A2a receptor; D1R, dopamine D1 receptor.

Fluor 488-coupled anti-chicken (1:500, Thermo Fisher Scientific Cat# A-11039, RRID:AB_2534096) antibodies. Sections were rinsed for 10 minutes (twice) in TBS and twice in Tris-buffer (1 M, pH 7.5) before mounting in DPX (Sigma-Aldrich). Confocal microscopy and image analysis were carried out at the Montpellier RIO Imaging Facility. All images covering the TS were single confocal sections acquired using sequential laser scanning confocal microscopy (Leica SP8) and stitched together as a single image. Double-labeled images from each region of interest were also single confocal sections obtained using sequential laser scanning confocal microscopy (Leica SP8 and Zeiss LSM780). eGFP-immunopositive cells were pseudocolored cyan and other immunoreactive markers were pseudocolored magenta or yellow as indicated in figures. Images used for quantification were all single confocal sections. Cells were considered positive for a given marker (eGFP and P-ERK) only when the nucleus was clearly visible. The D2R/A2AR-lacking, intermingled and the AST areas were identified using *Drd2-eGFP* mice or the anti-A2aR antibody. Quantification was performed manually using the cell counter plugin of the ImageJ software (RRID:SCR_003070) taking as standard reference a fixed threshold of fluorescence. Figures represent the number of neurons averaged from 3-4 adjacent sections.

### Brain slices preparation

Experiments were performed with 8-11 days old C57Bl/6J mice. Mice were housed under standardized conditions with a 12 hrs light/dark cycle, stable temperature (22 ± 1°C), controlled humidity (55 ± 10%) and food and water *ad libitum*. All animal procedures were performed in accordance with the Sorbonne University animal care committee regulations and the French Ministry of Agriculture and Forestry guidelines for handling animals. The protocol has been described previously in more details (Yapo *et al.* 2017). Briefly, 300 µm-thick coronal brain slices of were cut with a VT1200S microtome (Leica, RRID:SCR_008960). Slices were prepared in an ice-cold solution of the following composition: 125 mM NaCl, 0.4 mM CaCl_2_, 1 mM MgCl_2_, 1.25 mM NaH_2_PO_4_, 26 mM NaHCO_3_, 20 mM glucose, 2.5 mM KCl, 5 mM sodium pyruvate and 1 mM kynurenic acid, saturated with 5% CO_2_ and 95% O_2_. The slices were incubated in this solution for 30 minutes and then placed on a Millicell-CM membrane (Millipore) in culture medium (50% Minimum Essential Medium, 50% Hanks’ Balanced Salt Solution, 5.5 g/L glucose, penicillin-streptomycin, Invitrogen). We used the Sindbis virus as a vector to induce expression of the AKAR3 biosensors after overnight incubation. The coding sequences of AKAR3 (Allen and Zhang 2006) were inserted into the viral vector pSinRep5 (Invitrogen, San Diego, CA). This viral vector was added on the brain slices (∼5 × 10^5^ particles per slice), and the infected slices were incubated overnight at 35°C under an atmosphere containing 5% CO_2_. Before the experiment, slices were incubated for 30 min in the recording solution (125 mM NaCl, 2 mM CaCl_2_, 1 mM MgCl_2_, 1.25 mM NaH_2_PO_4_, 26 mM NaHCO_3_, 20 mM glucose, 2.5 mM KCl and 5 mM sodium pyruvate saturated with 5% CO_2_ and 95% O_2_). Recordings were performed with a continuous perfusion of the same solution at 32°C.

### Biosensor imaging and data analysis

Live imaging was performed on brain slices, which expressed the AKAR3 biosensor (Polito *et al.* 2014). Slices were kept at room temperature in the recording solution saturated with 5% CO_2_ and 95% O_2_ for around 30 minutes before imaging. The composition of the recording solution was the same as the slicing solution except that it contained no kynurenic acid and CaCl_2_ was 2 mM. The slice was imaged in a temperature controlled (32°C) recording chamber and continuously perfused at 2 ml/min using a peristaltic pump. The same perfusion system was used to apply drugs (SKF38393, Cat. #0922; CGS21680, Cat. #1063; Forskolin, Cat. #1099 purchased from Tocris, RRID:SCR_003689) onto the recorded slice. Wide field imaging was performed using Olympus BX50WI or BX51WI upright microscopes with a 40X 0.8 NA water immersion objective. The donor excitation was provided by an LED source (420 nm diode, Mightex) with a HC434/17 excitation filter (Semrock). The dichroic mirror was BS452 (Semrock). Two images were acquired for every data point by alternating filter for donor (HC479/40) and acceptor (HC542/50 both from Semrock) emission with a filter wheel (Sutter Instruments). Images were acquired using an ORCA-AG camera (Hamamatsu). Image acquisition was controlled using iVision software (Biovision, RRID:SCR_005057).

Data analysis was done with custom routines written in IGOR Pro (Wavemetric, RRID:SCR_000325) following the previously described protocol (Polito *et al.* 2014; Yapo *et al.* 2017). FRET changes were quantified using ratio of images from donor and acceptor channels for each data point. Corrections for non-uniform illumination, movement and channel mis-registration were done prior to ratio calculation. No corrections for bleed-through or cross-excitation were applied. The ratio value for a region-of-interest (ROI) was calculated by averaging all the individual pixels included in that ROI, weighted by their intensity. Unresponsive cells to either D1R or A2aR agonists were excluded.

### Statistical analysis

All statistical comparisons were performed with two-sided tests in Prism 6 (GraphPad Software, La Jolla, CA, USA, RRID:SCR_002798). The distribution of data was determined with Shapiro-Wilk normality test. No sample size calculations were performed. To determine outliers in every experimental group we performed the Grubbs’ test in Prism (RRID:SCR_002798). No outliers were identified. All the data were analyzed using either Student *t-test* with equal variances or One-way ANOVA. In all cases, significance threshold was automatically set at *p* < 0.05 (Prism 6). The One-way ANOVA analysis, where treatment was the independent variable, was followed by Bonferroni *post hoc* test for specific comparisons only when overall ANOVA revealed a significant difference (at least *p* < 0.05) among groups. All the values and statistical analyses have been reported in details in Supplemental Tables 1 to 12.

## Results

### Distribution of D1R- and D2R-SPNs in the TS is unique

*Drd2*-eGFP/*Drd1a*-tdTomato transgenic mice were used to characterize the distribution of D1R- and D2R-expressing SPNs in the TS. The analysis of eGFP (D2R-SPNs) and RFP (D1R-SPNs) expression revealed that, unlike a large portion of the dorsal striatum (Gangarossa *et al.* 2013b), the distribution of D1R- and D2R-positive cells in the TS was not random (Fig. 2a). At least three distinct territories were identified: *1*) a large strip adjacent to the GPe and devoid of D2R/A2aR-SPNs called hereafter the D2R/A2AR-lacking area, *2*) an intermingled area in which a random distribution of D1R- and D2R-SPNs was observed, and *3*) the amygdalostriatal transition area (AST) characterized by a high percentage of SPNs expressing both D1R and D2R (∼33%) (Fig. 2a, b). To confirm this non-random distribution, TS sections of *Drd2*-eGFP mice were used to analyze the endogenous eGFP fluorescence in combination with specific markers of D2R- and D1R-SPNs. D2R and A2aR antibodies, that stain D2R-SPNs dendrites, confirmed the existence of the D2R/A2AR-lacking area. Conversely, the strong D1R labeling confirmed that this subterritory of the TS was exclusively composed of D1R-SPNs (Fig. 2c).

**Figure 2:**
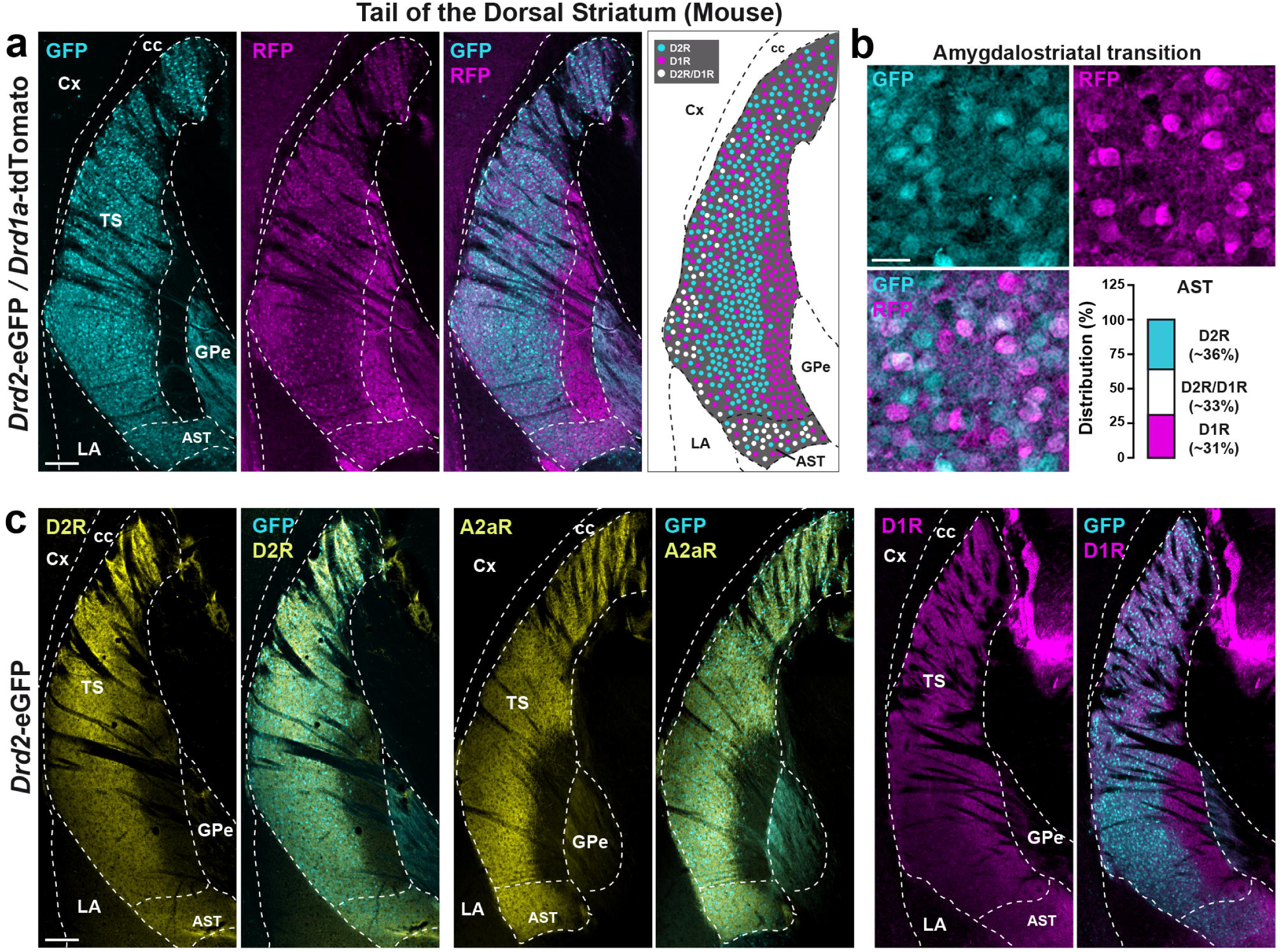
Distribution of D1R- and D2R-SPNs in the tail of the striatum. (a) RFP (magenta) and GFP (cyan) immunofluorescence in the TS of *Drd2-*eGFP*/Drd1a-* dtTomato mice. Scale bar, 200 µm. Schematic cartoon illustrating the distribution of GFP-, RFP- and GFP/RFP-expressing neurons. (b) High magnification of the AST of *Drd2-*eGFP*/Drd1a-*dtTomato mice. Scale bar, 20 µm. Distribution expressed as a percentage of neurons expressing GFP (D2R), RFP (D1R) or both GFP/RFP (D2R/D1R). (c) Double immunofluorescence of D2R, A2aR (yellow), D1R (magenta) and GFP (Cyan) in the TS of *Drd2-*eGFP mice. Scale bar, 200 µm. Cx: Cortex; cc: corpus callosum; GPe: external globus pallidus; AST: amygdalostriatal transition; LA: lateral amygdala; TS: tail of the striatum.

The peculiar distribution of D1R- and D2R/A2aR-SPNs in the TS led us to investigate the responsiveness of SPNs located in the D2R/A2aR-lacking and the AST areas in response to the stimulation of D1R and A2aR. In the rostral part of the dorsal striatum, stimulation of both receptors is known to yield an increase in cAMP levels and consequent activation of PKA in D1R- and D2R/A2aR-SPNs, respectively (Beaulieu and Gainetdinov 2011; Yapo *et al.* 2017). To address this issue, we performed *ex vivo* real-time PKA imaging in mouse TS slices by using the genetically encoded biosensor, AKAR3 (Yapo *et al.* 2017). In the D2R/A2aR-lacking area, stimulation of D1R with a saturating dose of SKF38393 (1 µM) yielded a large, but sub-maximal, increase in PKA signals in AKAR3-expressing neurons (Fig. 3a). In contrast, the application of the A2aR agonist, CGS21680 (1 µM), failed to further increase PKA signals. At the end of the experiment, addition of Forskolin (Fsk, 12.5 µM), an activator of adenylyl cyclase, maximally increased the F535/F480 emission ratio (Fig. 3a) (Castro *et al.* 2013). On average, CGS21680 had no effect on the amplitude of the PKA response induced by SKF38393 in the D2R/A2AR-lacking area (67 ± 3% vs 65 ± 2%, of the maximal Fsk response, respectively) (Fig. 3b). On the other hand, in the AST area, the amplitude of PKA responses produced by SKF38393 (62 ± 3% of the maximal Fsk response) was further increased in response to A2aR activation (94 ± 1%) (Fig. 3c, d). These results show that the D2R/A2AR-lacking area expresses functional D1 receptors and lacks A2A receptors, whereas both receptors are functional in the AST region. Together, these results confirm that SPNs of the D2R/A2AR-lacking area only respond to D1R agonist, whereas the AST responds to both D1R and A2aR activation.

**Figure 3:**
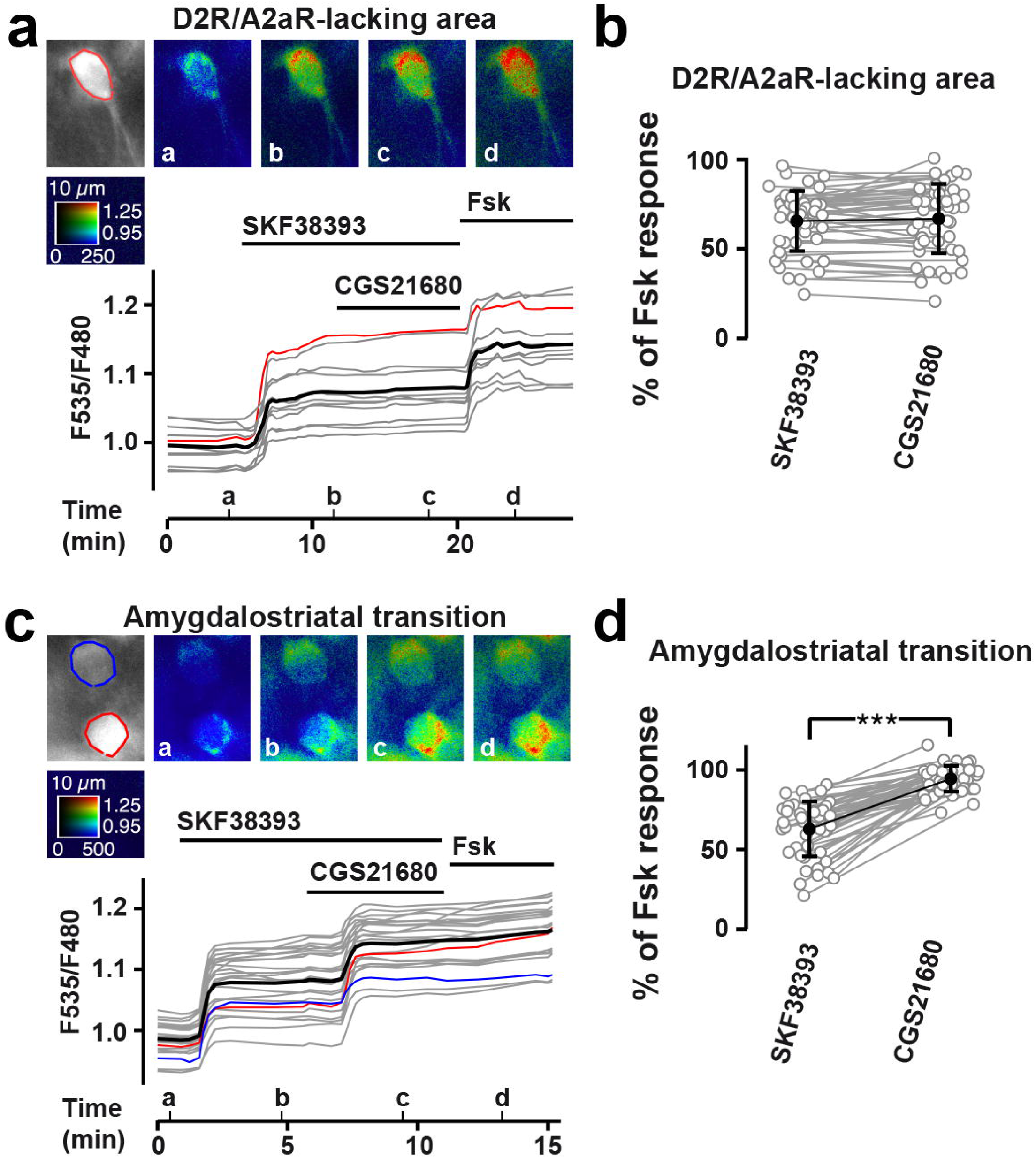
PKA imaging following activation of D1R and A2aR in the tail of the striatum. Wide-field imaging of mouse striatal brain slices expressing AKAR3 biosensor in D2R/A2aR-lacking (a, b) and in the AST (c, d) areas. (a, c) Raw fluorescence of the acceptor is displayed in gray and acceptor/donor fluorescence is displayed in pseudo-color, coded from blue (low F535/F480 ratio, ie low PKA signal) to red (high PKA signal). Ratio images (a-d) correspond to the time points indicated in the bottom graph. Changes in ratio were measured during time and each trace indicates the F535/F480 emission ratio measured on the region of interest of corresponding color shown on the gray image. Traces in gray correspond to regions of interest outside the selected region of the field. The thick black trace represents the average. Horizontal bars indicate the bath application of SKF38393 (1µM), CGS21680 (1µM) and forskolin (Fsk, 12.5 µM). (b) Amplitude of the F535/F480 emission ratio triggered by SKF38393 (SKF, n_neurons_ = 48 from 3 mice) and CGS21680 (CGS, n_neurons_ = 48 from 3 mice) in the D2R/A2aR-lacking area. (d) Amplitude of the F535/F480 emission ratio triggered by SKF38393 (SKF, n_neurons_ = 47 from 3 mice) and CGS21680 (CGS, n_neurons_ = 47 from 3 mice) in the AST. Mean ± sem is represented by the filled circles. CGS21680 vs SKF38393 *** p < 0.001. AST: amygdalostriatal transition. For detailed statistics see Supplemental Table 1.

### Distribution of D1R- and D2R-SPNs in the TS is evolutionarily conserved

We next wondered whether a similar organization of the TS might be observed in distinct species of the Muridae family. To explore this issue, we examined the presence of the D2R/A2AR-lacking area in different wild mouse species of the genus *Mus*. This genus comprises several species subdivided into four subgenera encompassing about 7 million years of divergence (Chevret and Dobigny 2005; Veyrunes *et al.* 2006) (Fig. 4a). The lack of A2aR immunoreactivity and the strong expression of DARPP-32, a highly conserved marker of all SPNs, were used here to define a molecular fingerprint of the D2R/A2AR-lacking area (Gangarossa *et al.* 2013b). Using this combination of antibodies, we observed that the D2R/A2AR-lacking area was present in all the species examined which included 3 subspecies of the *Mus musculus*, 4 species of *Subgenus Mus* and 2 species of two of the three other *Subgenera* (Fig. 4a-c). A similar organization of the caudal TS was also identified in *Meriones unguiculatus* (Mongolian gerbil) (Fig. 4d) and *Rattus norvegicus* (Fig. 4e). Taken together, our data reveal that the distribution of D1R- and D2R-SPNs in the TS is evolutionarily conserved.

**Figure 4:**
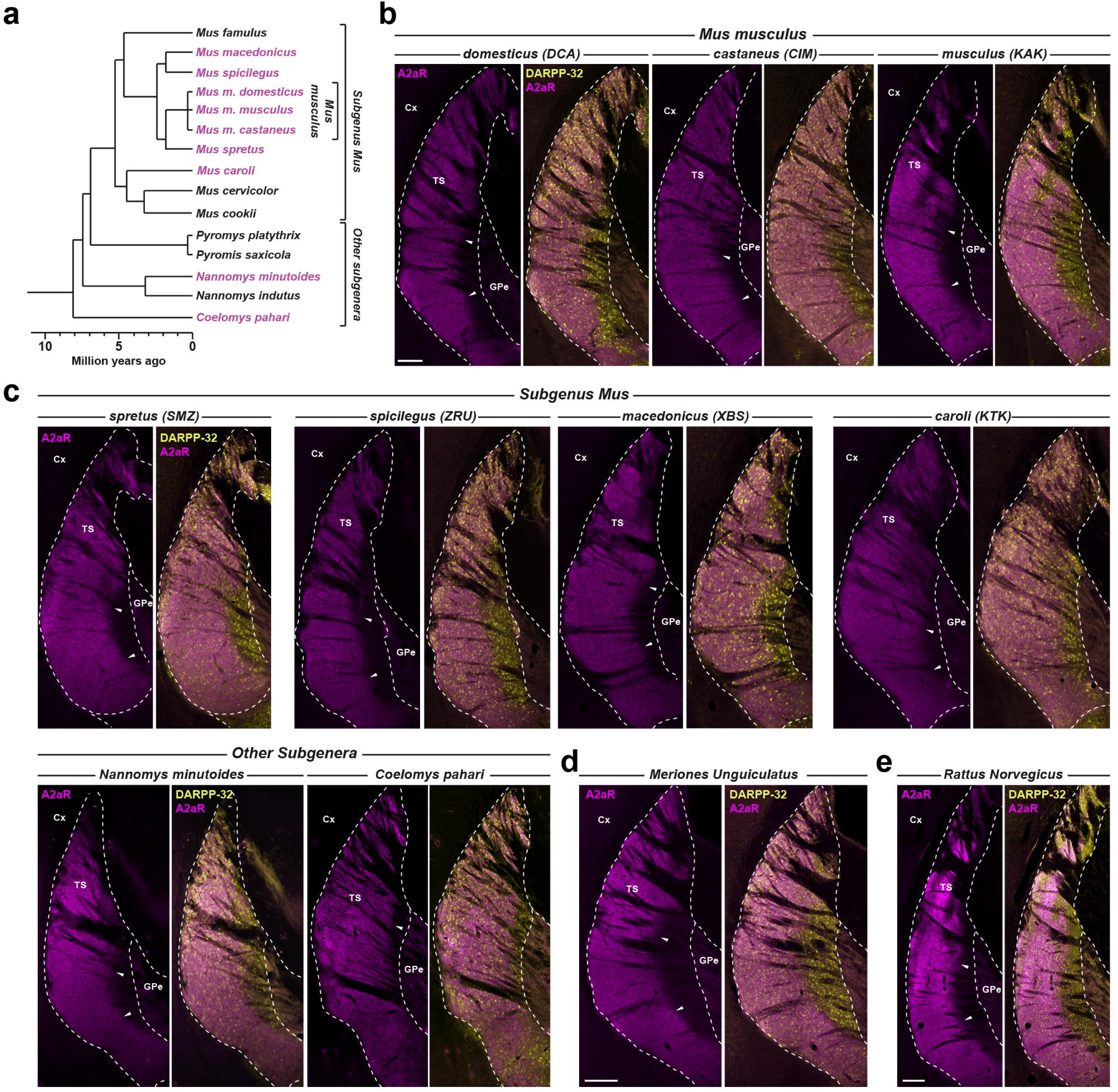
The D2R/A2aR-lacking area is evolutionarily conserved. (a) Simplified phylogenetic tree and divergence time estimates of the *Mus* species group, according to (Chevret and Dobigny 2005; Veyrunes *et al.* 2006). Species identified in magenta have been used in the present study. (b-e) Double immunofluorescence of A2aR (magenta) and DARPP-32 (yellow) in the TS of (b) 3 subspecies of the *Mus musculus* [*Mus m. domesticus* (DCA), the *Mus m. castaneus* (CIM) and the *Mus musculus* (KAK), c) 4 species of *Subgenus Mus* [*Mus spretus* (SMZ), *Mus spicilegus* (ZRU), *Mus macedonicus* (XBS) and *Mus caroli* (KTK)] and 2 species from other *Subgenera* (*Nannomys minutoides* and *Coelomys pahari*). Scale bar, 250 µm, (d) in *Meriones unguiculatus* (Mongolian gerbil). Scale bar, 200 µm, and (e) in *Rattus norvegicus.* Scale bar, 250 µm. White arrowheads delineate the boundaries between the intermingled area and D2R/A2aR-lacking area. Cx: Cortex; GPe: external globus pallidus; TS: tail of the striatum.

### Regulation of ERK activation in the TS by aversive signals

Avoidance behaviors have been associated with the activation of the TS (Menegas *et al.* 2018). To determine the impact of aversive stimuli on the TS, we monitored ERK activation by quantifying the number of phosphorylated-ERK (P-ERK) immunoreactive SPNs in the three territories of the TS: the intermingled area, the D2R/A2AR-lacking area and the AST (Fig. 5, Supplemental Fig. 1-3 and Table 3).

**Figure 5:**
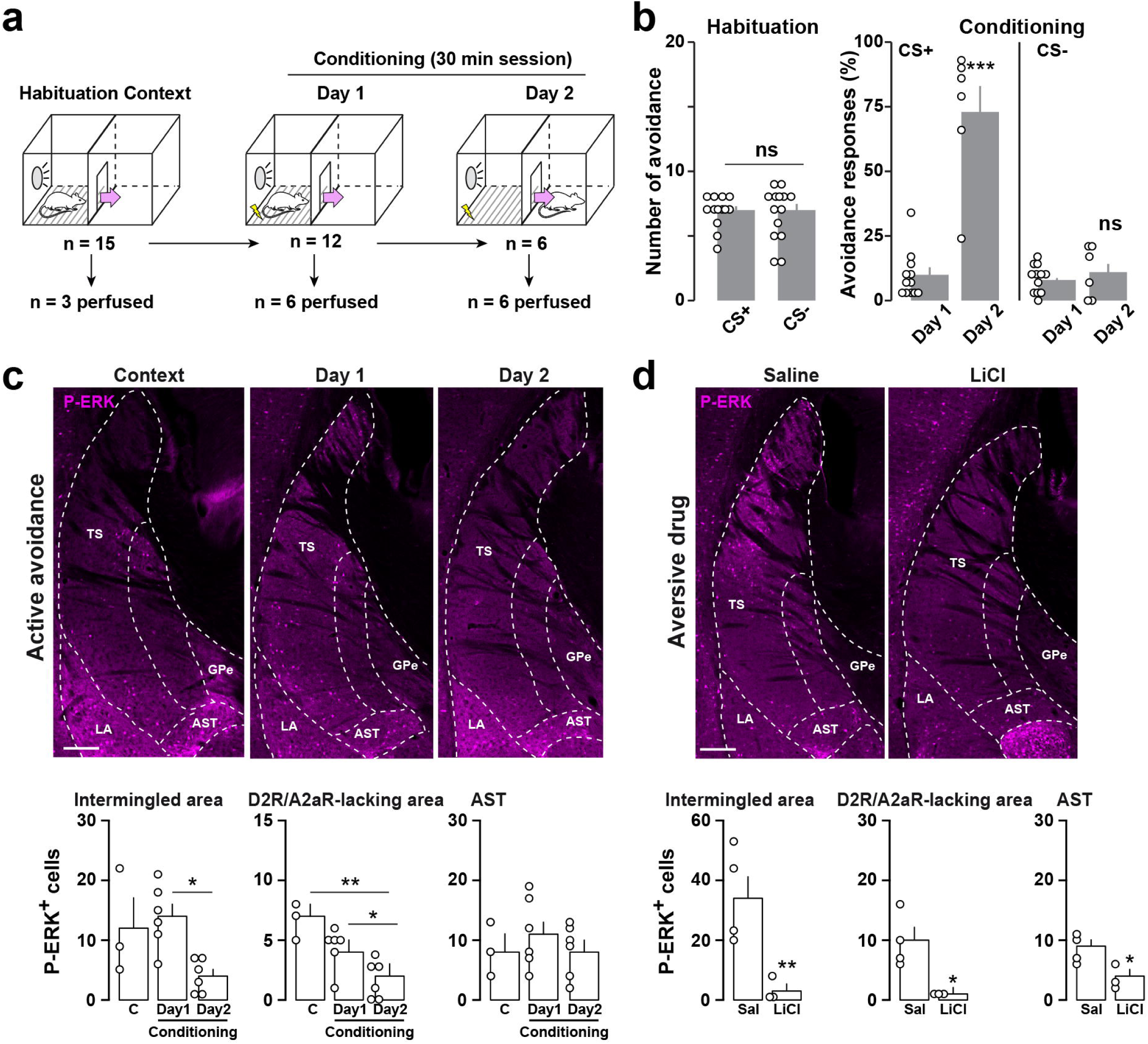
Regulation of ERK phosphorylation in the TS by aversive signals. (a) Drawing indicates the three phases of the active avoidance paradigm (habituation to context, conditioning Day 1 and conditioning Day 2) and the repartition of mice. (b) Active avoidance responses during the habituation (mice were only exposed to the shuttle box and the CS) and during the first and second session of conditioning (Day 1 and Day 2). Experimental groups: mice exposed to the context (n = 15), conditioned Day 1 (n = 12) and conditioned Day 2 (n = 6). Data are presented as means ± sem. *** p < 0.001 CS^+^ Day2 vs CS^+^ Day1. (c) P-ERK immunoreactivity (magenta) in the TS of C57Bl/6J mice exposed to active avoidance. Scale bar, 200 µm. Quantification of P-ERK-immunoreactive neurons in the intermingled, the D2R/A2aR-lacking and the AST areas of mice exposed to the context (n = 3) or conditioned D1 (n = 6) or D2 (n = 6) days. Data are presented as means ± sem. * p < 0.05, ** p < 0.01. (d) P-ERK immunoreactivity (magenta) in the TS of C57Bl/6J mice 15 min after saline or LiCl (150 mg/kg) administration. Scale bar, 200 µm. Quantification of P-ERK-immunoreactive neurons in the intermingled, the D2R/A2aR-lacking and the AST areas of mice injected with saline (n = 4) or LiCl (n = 3). Data are presented as means ± sem. * p < 0.05, ** p < 0.01, TS: tail of the striatum; GPe: external globus pallidus; AST: amygdalostriatal transition; LA: lateral amygdala. For detailed statistics see Supplemental Table 2.

**Table 3.** Regulation of ERK phosphorylation in the TS by aversive signals.

| **Models/Drugs**<br>(*n* = number of mice) | **Average number of P-ERK-positive cells** |  |  |  |
| --- | --- | --- | --- | --- |
|  | **Total** | **Intermingled** | **D2R/A2aR-lacking** | **AST** |
| **3.1 Resident/Intruder** |  |  |  |  |
| *Resident (n = 3)* | 50 ± 5 | 32 ± 3 | 9 ± 1 | 8 ± 1 |
| *Intruder (n = 3)* | 56 ± 3 | 28 ± 3 | 14 ± 2 | 14 ± 2 |
| **3.2 Natural stressors** |  |  |  |  |
| *Mouse bedding (n = 3)* | 71 ± 6 | 38 ± 6 | 10 ± 3 | 15 ± 1 |
| *Rat bedding (n = 3)* | 66 ± 14 | 43 ± 10 | 13 ± 3 | 10 ± 2 |
| *Water (n = 4)* | 38 ± 10 | 25 ± 7 | 5 ± 2 | 9 ± 2 |
| *Fox urine (TMT) (n = 3)* | 52 ± 3 | 28 ± 3 | 7 ± 1 | 17 ± 4 |
| **3.3 Cued fear conditioning** |  |  |  |  |
| *Context (n = 3)* | 29 ± 9 | 16 ± 5 | 1 ± 1 | 12 ± 4 |
| *CS (n = 6)* | 33 ± 13 | 16 ± 9 | 2 ± 1 | 15 ± 4 |
| *US (n = 6)* | 53 ± 6 | 32 ± 5 | 3 ± 1 | 17 ± 1 |
| *CS-US paired (n = 6)* | 41 ± 2 | 23 ± 3 | 4 ± 1 | 14 ± 1 |
| *CS-US unpaired (n = 3)* | 57 ± 9 | 36 ± 7 | 5 ± 1 | 16 ± 3 |

For detailed statistics see Supplemental Table 7.
CS, conditioned stimulus; US, unconditioned stimulus; TMT, 2,3,5-trimethyl-3-thiazoline

We first analyzed the level of P-ERK in mice that underwent the resident-intruder assay. The resident and intruder mice were perfused ten minutes after the unfamiliar mouse was placed inside the resident’s cage. No significant differences were found in the number of P-ERK-positive neurons in the TS (Supplemental Fig. 2a and Table 3). We then tested whether exposure to predator odors was able to modulate P-ERK in the TS. Although mice exposed to rat bedding or 2,3,5-Trimethyl-3-thiazoline (TMT), a component of fox odor, showed strong defensive reactions, ERK activation remained unchanged in all territories of the TS when compared to control mice (Supplemental Fig. 2b and Table 3). Taken together, these data suggest that the TS is not recruited by aversive stimuli triggering innate avoidance.

We therefore assessed whether mice exposed to mild footshocks, either unpredictable (US) or signaled by a salient auditory stimulus (arousing but non-nociceptive) (CS-US pairing), displayed an altered P-ERK level in the caudal TS (Supplemental Fig. 3a, b and Table 3). No changes were observed in the TS of mice undergoing five CS-US paired or unpaired compared to those only exposed to the CS, the US or the context (Supplemental Fig. 3b and Table 3). We then monitored ERK phosphorylation in the TS of mice that underwent active avoidance paradigm. Mice that learned to avoid mild footshocks during an interval signaled by the presentation of a CS (salient auditory stimulus) displayed a mild, but significant, reduction in the number of P-ERK-immunoreactive neurons in the TS, in particular in the intermingled and the D2R/A2AR-lacking areas (Fig. 5a-c). We finally thought to determine whether prolonged aversive states might alter the levels of P-ERK in the TS. The administration of lithium chloride (LiCl) (150 mg/kg), a compound known to cause nausea and visceral malaise (Yamamoto and Ueji 2011), blunted ERK activation in all TS subterritories (Fig. 5d).

Because the effects induced by the administration of LiCl can also be perceived as potent anorexigenic signals, we investigated whether SPNs of the TS were activated during food deprivation (fasting) and satiation (refeeding). Indeed, food deprivation triggered a significant reduction of the body weight and an increase in food intake during refeeding schedules (Supplemental Fig. 4). However, no major differences were observed in the number of P-ERK immunoreactive neurons between mice having *ad libitum* access to chow pellets and mice being fasted for 12 hours followed or not by 1 or 6 cycles of re-feeding schedule (Table 4 and Supplemental Fig. 4). Similarly, ERK activation was unchanged in the TS of mice exposed to high-fat high-sugar (HFHS) diet during 1h/day (Table 4 and Supplemental Fig. 4). Together, these data suggest that SPNs of the TS are not activated by signals associated with fasting, satiety or food palatability.

**Table 4.** Regulation of ERK phosphorylation in the TS by satiety and palatable signals.

| **Models**<br>(*n* = number of mice) | **Average number of P-ERK-positive cells** |  |  |  |
| --- | --- | --- | --- | --- |
|  | **Total** | **Intermingled** | **D2R/A2aR-lacking** | **AST** |
| **4.1 Food intake** |  |  |  |  |
| *Control* (*n* = 4) | 39 ± 5 | 29 ± 4 | 2 ± 1 | 8 ± 1 |
| *Fast* (*n* = 3) | 23 ± 6 | 15 ± 5 | 2 ± 1 | 6 ± 1 |
| *Fast/re-fed*<sub>1cycle</sub> (*n* = 4) | 32 ± 2 | 21 ± 1 | 2 ± 1 | 9 ± 1 |
| *Fast/re-fed*<sub>6cycles</sub> (*n* = 4) | 49 ± 9 | 32 ± 7 | 4 ± 1 | 13 ± 3 |
| *HFHS*<sub>binge</sub> (*n* = 4) | 36 ± 1 | 22 ± 2 | 4 ± 1 | 10 ± 1 |

For detailed statistics see Supplemental Table 8.
HFHS, high-fat high-sugar diet

### Psychostimulant drugs promote ERK activation in the TS

Acute d-amphetamine administration activates ERK in striatal and accumbal D1R-SPNs (Valjent *et al.* 2005; Gerfen *et al.* 2008; Gangarossa *et al.* 2013a). We therefore analyzed whether the same treatment was also able to promote ERK phosphorylation in the TS. Immunofluorescence analysis revealed that a single administration of d-amphetamine (10 mg/kg) increased the number of P-ERK-positive neurons in the intermingled and D2R/A2AR-lacking areas of the TS (Supplemental Fig. 5a, b). Of note, only a mild increase was observed in the AST (Supplemental Fig. 5a, b). To clarify in which striatal cell-type ERK activation occurred, *Drd2*-eGFP mice were treated with d-amphetamine and the distribution of P-ERK-positive neurons in D2R-positive and D2R-negative SPNs was analyzed. In line with the observations made in the rostral part of the dorsal striatum (DS) (Valjent *et al.* 2005; Gerfen *et al.* 2008), d-amphetamine-induced ERK phosphorylation in the TS was confined in D2R-negative SPNs, which most likely correspond to D1R-SPNs (Fig. 6a, b). This increase in ERK phosphorylation was rapid and transient since it declined 30 min after the administration of d-amphetamine (Fig. 6b, Supplemental Fig. 5b). In the rostral DS, ERK activation induced by d-amphetamine is preferentially concentrated in patches (Biever *et al.* 2015). We therefore thought to determine whether P-ERK-immunoreactive neurons of the caudal TS were also distributed within patches. To do so, we used *Nr4a1*-eGFP mice, a mouse line allowing the delineation of the patch/matrix compartmentalization (Davis and Puhl 2011). The analysis of eGFP expression revealed that eGFP-positive cells were mainly located in the intermingled area whereas only few scattered neurons were detected in the D2R/A2AR-lacking area and AST (Fig. 6c). When d-amphetamine was administrated to *Nr4a1*-eGFP mice, we observed that the increased P-ERK-positive cells occurred preferentially in eGFP-poor zones and was restricted to *Nr4a1*-eGFP-negative neurons suggesting that in the TS, ERK activation was not enriched in patch compartments (Fig. 6c, d).

**Figure 6:**
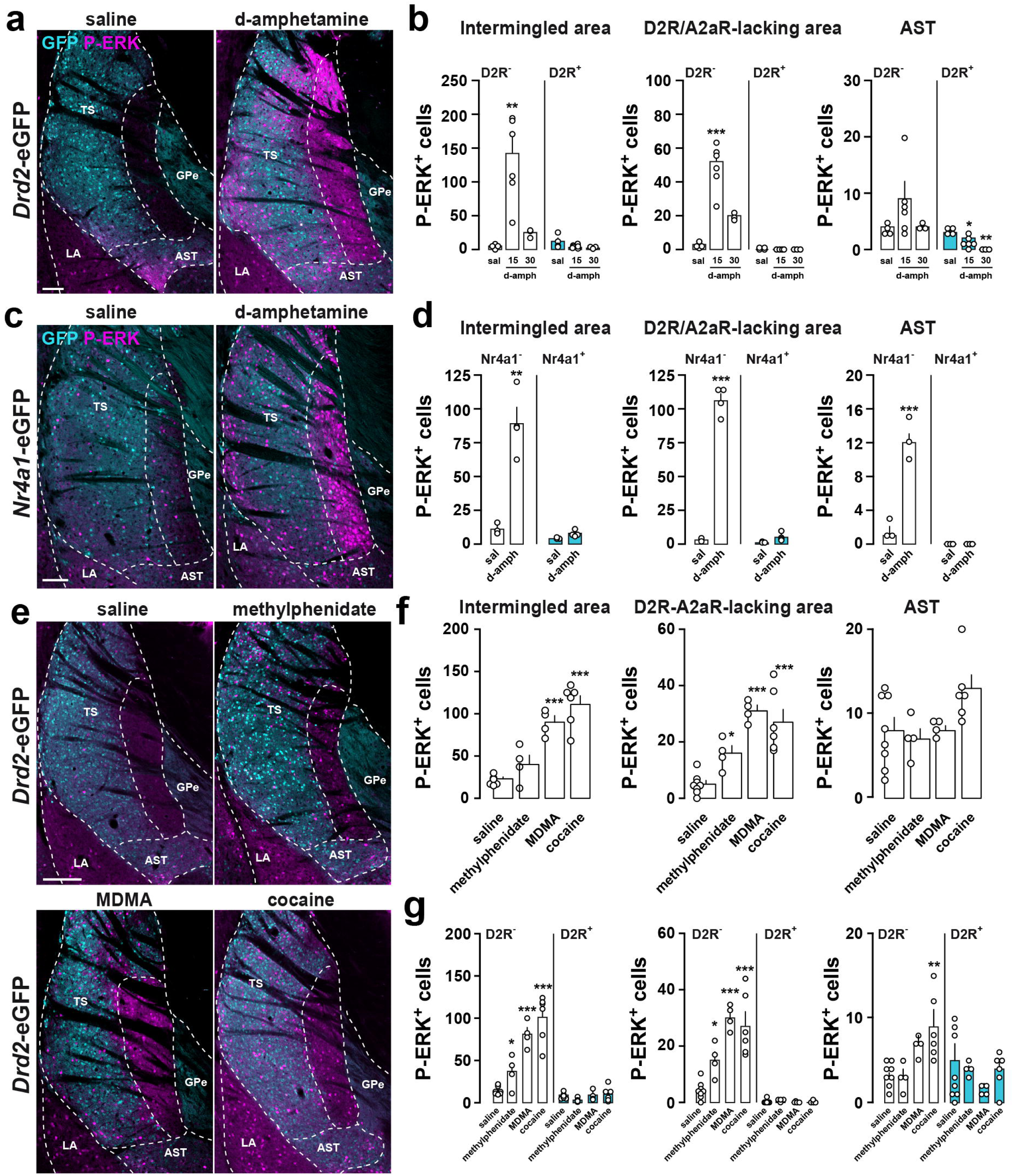
Psychostimulants increase ERK phosphorylation in the TS. (a) P-ERK (magenta) and GFP (cyan) immunoreactivity in the TS of *Drd2-*eGFP mice 15 min after saline or d-amphetamine (10 mg/kg) administration. Scale bar, 100 µm. (b) Quantification of P-ERK-immunoreactive neurons among eGFP-positive (D2R+) or eGFP-negative (D2R-) cells in the intermingled, the D2R/A2aR-lacking and the AST areas of *Drd2-*eGFP mice treated saline (n = 4) or d-amphetamine and perfused 15 (n = 6) or 30 min (n = 3) after injection. Data are presented as means ± sem. ** p < 0.01, *** p < 0.001 sal vs d-amph. (c) P-ERK (magenta) and GFP (cyan) immunoreactivity in the TS of *Nr4a1-*eGFP mice 15 min after saline (n = 3) or d-amphetamine (10 mg/kg, n = 4) administration. Scale bar, 100 µm. (d) Same quantifications as in (b) performed in *Nr4a1-*eGFP mice. Data are presented as means ± sem. ** p < 0.01, *** p < 0.001 sal vs d-amph. (e) P-ERK (magenta) and GFP (cyan) immunoreactivity in the TS of *Drd2-*eGFP mice 15 min after saline (n = 8), methylphenidate (15 mg/kg, n = 4), MDMA (10 mg/kg, n = 4) or cocaine (15 mg/kg, n = 6) administration. Scale bar, 200 µm. (f) Quantification of P-ERK-immunoreactive neurons in the intermingled, the D2R/A2aR-lacking and the AST areas of mice treated with saline (n = 8), methylphenidate (15 mg/kg, n = 4), MDMA (10 mg/kg, n = 4) or cocaine (15 mg/kg, n = 6). Data are presented as means ± sem. * p < 0.05, ** p < 0.01, *** p < 0.001 sal vs drugs. (g) Quantification of P-ERK-immunoreactive neurons among eGFP-positive (D2R+) or eGFP-negative (D2R-) cells in the intermingled, the D2R/A2aR-lacking and the AST areas of *Drd2-*eGFP mice treated saline (n = 8), methylphenidate (15 mg/kg, n = 4), MDMA (10 mg/kg, n = 4) or cocaine (15 mg/kg, n = 6) administration and perfused 15 after injection. Data are presented as means ± sem. * p < 0.05, ** p < 0.01, *** p < 0.001 sal vs drugs. GPe: external globus pallidus; AST: amygdalostriatal transition; LA: lateral amygdala; TS: tail of the striatum; sal: saline; d-amph: d-amphetamine. See also Supplemental Fig. 5. For detailed statistics see Supplemental Table 3 and 12.

Finally, we tested whether other psychostimulant drugs shared the same pattern of ERK activation in the TS. As shown in Figure 6e-g, a single injection of methylphenidate (15 mg/kg), MDMA (10 mg/kg) or cocaine (15 mg/kg) increased ERK phosphorylation in the D2R/A2AR-lacking and the intermingled areas in which it was restricted to D2R-negative SPNs. Together, these data revealed that in the TS, D1R-SPNs of the intermingled and D2R/A2AR-lacking areas are activated by psychostimulant drugs.

In addition to its stimulant properties, MDMA is also known to distort sensory and time perception, two features shared by hallucinogenic drugs (Starr *et al.* 2008; Litjens *et al.* 2014). We therefore tested whether other hallucinogen drugs were able to activate SPNs of the TS. The administration of the psychedelic compound, DOI (10 mg/kg), a substituted amphetamine without stimulant effects, failed to induce ERK activation in the TS (Table 5). On the other hand, PCP (3 mg/kg), a NMDA receptor blocker causing hallucinations, reduced P-ERK in the caudal TS (Table 5). This decrease was even more pronounced after the administration of dizocilpine (0.3 mg/kg), another NMDA blocker that also acts as a potent anti-convulsant (Table 5). Because convulsant drugs also trigger ERK activation (Gangarossa *et al.* 2011; Berkeley *et al.* 2002), we examined whether such pharmacological agents were able to regulate ERK phosphorylation in the TS. Indeed, acute administration of PTZ (60 mg/kg) or kainate (20 mg/kg) increased the number of P-ERK-positive neurons exclusively in the AST area (Table 5). Together, these results indicate that, at the level of the TS, distinct classes of pharmacological agents differentially recruit D1R- and D2R/A2aR-SPNs in distinct subsets of striatal territories.

**Table 5.** Regulation of ERK phosphorylation in the TS by hallucinogens and convulsants.

| **Drugs**<br>(*n* = number of mice) | **Average number of P-ERK-positive cells** |  |  |  |
| --- | --- | --- | --- | --- |
|  | **Total** | **Intermingled** | **D2R/A2aR-lacking** | **AST** |
| **5.1 Hallucinogens** |  |  |  |  |
| *Control* (*n* = 4) | 47 ± 5 | 29 ± 5 | 11 ± 1 | 7 ± 1 |
| *DOI* (*n* = 4) | 47 ± 8 | 32 ± 5 | 5 ± 1 | 10 ± 3 |
| *PCP* (*n* = 3) | 25 ± 6 | 11 ± 1 | 10 ± 4 | 4 ± 1 |
| *Dizocilpine* (*n* = 3) | 4 ± 1*** | 0 ± 0** | 1 ± 1** | 3 ± 1 |
| **5.2 Convulsants** |  |  |  |  |
| *Control* (*n* = 3) | 33 ± 5 | 21 ± 2 | 4 ± 2 | 8 ± 1 |
| *PTZ* (*n* = 3) | 72 ± 7 | 27 ± 2 | 7 ± 1 | 38 ± 7** |
| *Kainate* (*n* = 3) | 69 ± 15 | 34 ± 15 | 10 ± 4 | 25 ± 2* |

For detailed statistics see Supplemental Table 9.
DOI, 2,5-Dimethoxy-4-iodoamphetamine; PCP, phencyclidine; PTZ, Pentylentetrazol.

### Increased ERK activation in the TS by selective DA reuptake inhibition

Psychostimulant drugs such as d-amphetamine, methylphenidate, cocaine and MDMA mediate their effects in part by acting on monoamines transporters (DAT, NET and SERT). We therefore examined whether the blockade of dopamine, noradrenaline or serotonin reuptake promoted ERK activation in the TS. Administration of GBR12783 (10 mg/kg), a DAT blocker, enhanced ERK phosphorylation preferentially in SPNs of the D2R/A2AR-lacking area and to a lesser extent in the intermingled area (Fig. 7a, b). In contrast, the antidepressant drugs desipramine (20 mg/kg) and fluoxetine (10 mg/kg), which inhibit the reuptake of NE and 5-HT respectively, failed to activate ERK (Fig. 7a, b). Together, these results show that blockade of DA reuptake is sufficient to trigger ERK activation in the TS.

**Figure 7:**
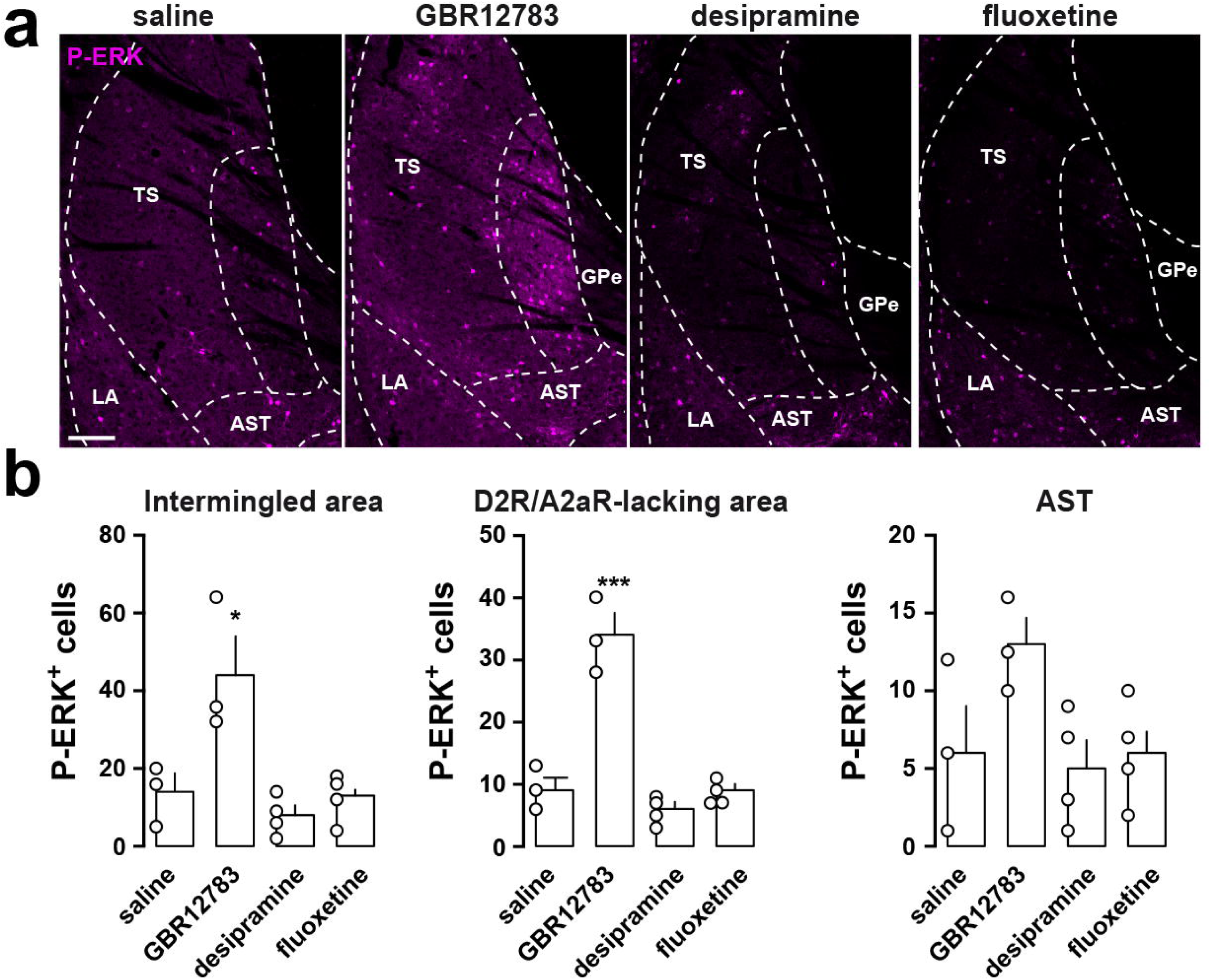
Dopamine reuptake inhibitor enhances ERK activation in the TS. (a) P-ERK (magenta) immunoreactivity in the TS of C57Bl/6J mice 15 min after saline (n = 3), GBR12783 (10 mg/kg, n = 3), desipramine (20 mg/kg, n = 4) or fluoxetine (10 mg/kg, n = 4) administration. Scale bar, 100 µm. (b) Quantification of P-ERK-immunoreactive neurons in the intermingled, the D2R/A2aR-lacking and the AST areas of mice treated with saline (n = 3), GBR12783 (10 mg/kg, n = 3), desipramine (20 mg/kg, n = 4) or fluoxetine (10 mg/kg, n = 4). Data are presented as means ± sem. * p < 0.05, *** p < 0.001 sal vs drugs. GPe: external globus pallidus; AST: amygdalostriatal transition; LA: lateral amygdala; TS: tail of the striatum. For detailed statistics see Supplemental Table 4.

### D1R activation is necessary and sufficient to activate ERK in the TS

In the rostral DS and the nucleus accumbens, ERK activation induced by d-amphetamine requires D1R stimulation (Valjent *et al.* 2005). Similarly, blockade of D1R by SCH23390 prevented d-amphetamine-induced ERK phosphorylation in the TS (Fig. 8a, b). However, contrasting results have been obtained regarding the ability of D1R to promote ERK phosphorylation in the rostral DS (Gerfen *et al.* 2002; Gangarossa *et al.* 2013c). To test the impact of D1R activation, mice were administrated either with SKF81297 (5 mg/kg) or SKF83822 (5 mg/kg), two agonists known to trigger D1R-dependent signaling through cAMP production (Kuroiwa *et al.* 2008). Both compounds produced a strong activation of ERK that resembled the one induced by d-amphetamine (Fig. 8c, d). In contrast, when we injected SKF83959 (5 mg/kg), a partial agonist at both D1R-mediated adenylate cyclase and β-arrestin recruitment (Lee *et al.* 2014; Jin *et al.* 2003), ERK phosphorylation was less pronounced (Fig. 8c, d). Together, these results indicate that D1R stimulation is sufficient to activate ERK in the TS.

**Figure 8:**
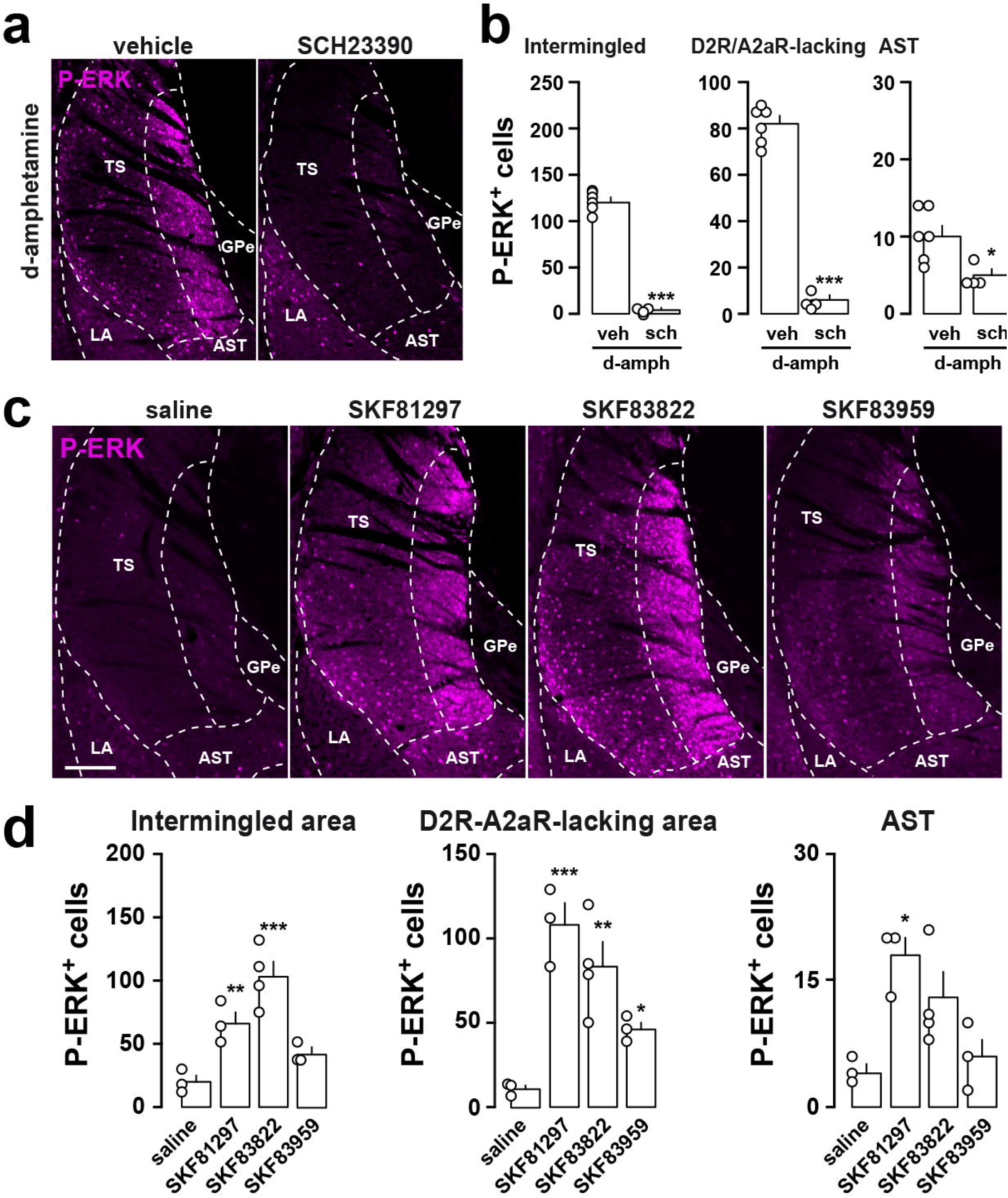
D1R-dependent ERK phosphorylation in the TS. (a) P-ERK (magenta) immunofluorescence in the TS of d-amphetamine-injected mice, in the presence (n = 4) or not (n = 6) of a D1R antagonist (sch) injected 15 min before d-amphetamine. Scale bar, 200 µm. (b) Quantification of P-ERK-immunoreactive neurons in the intermingled, the D2R/A2aR-lacking and the AST areas of mice treated d-amphetamine alone (n = 6) or with SCH23390 (n = 4). Data are presented as means ± sem. * p < 0.05, ** p < 0.01 veh/d-amph vs sch/d-amph. (c) P-ERK (magenta) immunofluorescence in the TS of mice injected with saline (n = 3), SKF81297 (5 mg/kg, n = 3), SKF83822 (5 mg/kg, n = 4) and SKF83959 (5 mg/kg, n = 3). Scale bar, 200 µm. (d) Quantification of P-ERK-immunoreactive neurons in the intermingled, the D2R/A2aR-lacking and the AST areas of mice treated with saline (n = 3), SKF81297 (5 mg/kg, n = 3), SKF83822 (5 mg/kg, n = 4) and SKF83959 (5 mg/kg, n = 3). Data are presented as means ± sem. * p < 0.05, ** p < 0.01, *** p < 0.001 saline vs SKF. GPe: external globus pallidus; AST: amygdalostriatal transition; LA: lateral amygdala; TS: tail of the striatum; sch: SCH23390; veh: vehicle; sal: saline; d-amph: d-amphetamine. For detailed statistics see Supplemental Table 5.

To determine whether cAMP production was necessary and sufficient to activate ERK in the TS, we artificially increased the cAMP levels by administering papaverine, a potent inhibitor of the striatal-enriched phosphodiesterase isoform PDE10A (Nishi *et al.* 2008). Papaverine (30 mg/kg) induced a robust ERK activation in the TS (Fig. 9a, b). However, this activation was restricted to the ventral part of the intermingled area and confined in D2R-SPNs (Fig. 9a-c). Because D1R- and D2R-SPNs also express PDE4, we also examined the impact of PDE4 inhibition. Administration of rolipram (10 mg/kg) significantly increased ERK phosphorylation in the AST area whereas little or no significant effects were observed in the intermingled and the D2R/A2aR-lacking areas, respectively (Fig. 9d, e). In this case, increased ERK phosphorylation was detected in D2R-negative SPNs (Fig. 9e, AST). Altogether, these results show that D1R stimulation or PDEs inhibition trigger distinct patterns of ERK phosphorylation in the TS.

**Figure 9:**
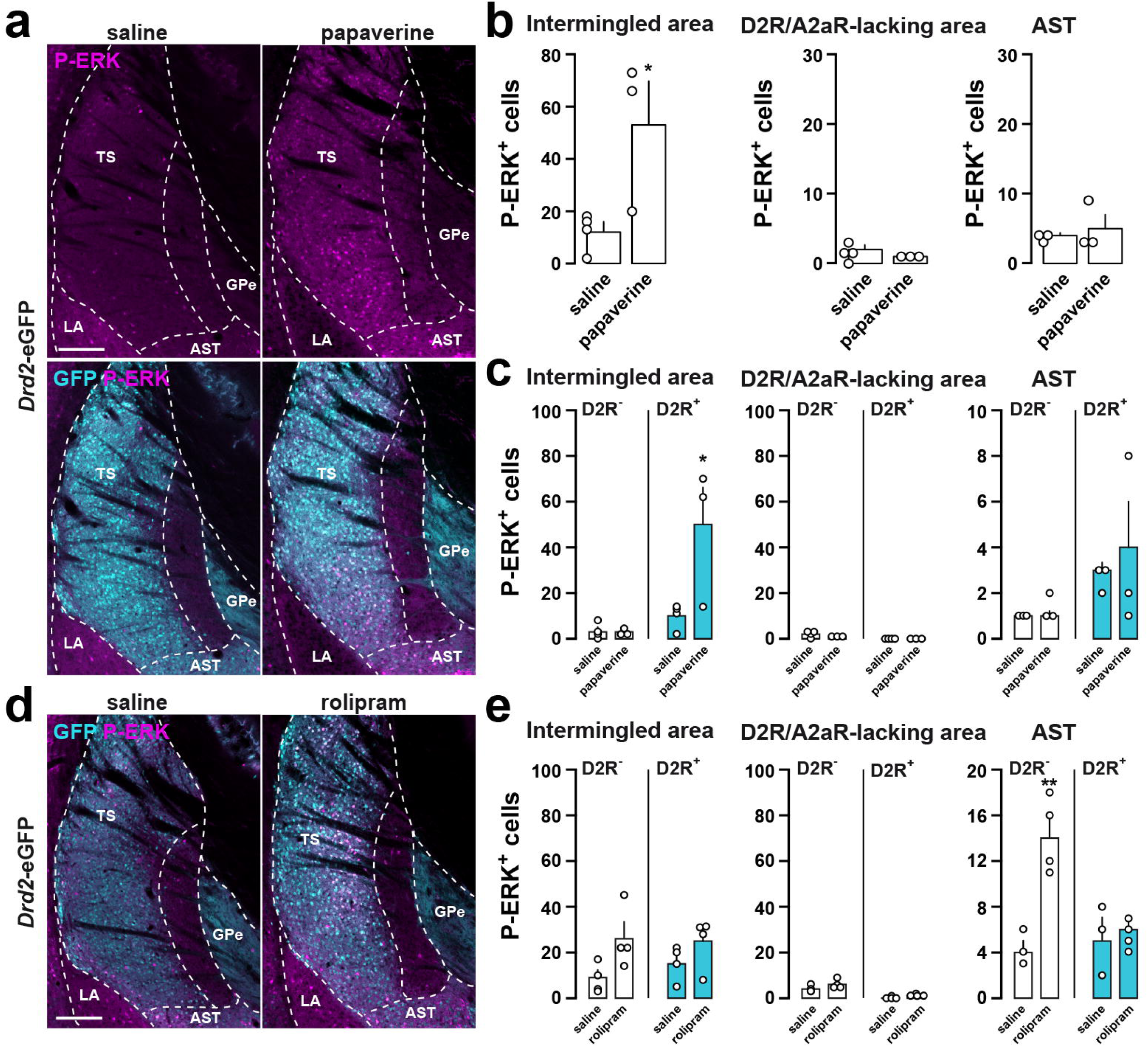
Distinct pattern of ERK phosphorylation in the TS following PDE inhibition. (a) P-ERK (magenta) and GFP (cyan) immunolabelling in the TS of *Drd2-* eGFP mice 15 min after saline (n = 4) or papaverine (30 mg/kg, n = 3) administration. Scale bar, 200 µm. (b) Quantification of P-ERK-immunoreactive neurons in the intermingled, the D2R/A2aR-lacking and the AST areas of mice treated with saline (n = 4) or papaverine (30 mg/kg, n = 3). Data are presented as means ± sem. (c) Quantification of P-ERK-immunoreactive neurons among eGFP-positive (D2R+) or eGFP-negative (D2R-) cells in the intermingled, the D2R/A2aR-lacking and the AST areas of *Drd2-*eGFP mice treated with saline (n = 4) or papaverine (30 mg/kg, n = 3). Data are presented as means ± sem. (d and e) Same analysis and quantification as in (c) except that mice were treated with saline (n = 4) or rolipram (10 mg/kg, n = 4). * p < 0.05, ** p < 0.01, *** p < 0.001 sal vs drugs. GPe: external globus pallidus; AST: amygdalostriatal transition; LA: lateral amygdala; TS: tail of the striatum. For detailed statistics see Supplemental Table 6.

## Discussion

Despite the renewed interest in the caudal part of the striatum (also referred to as the tail of the striatum, TS), its anatomo-functional organization remains poorly characterized. In this study, we provide evidence that D1R- and D2R-SPNs of the TS display a unique regional distribution whose anatomical features are to some extent evolutionarily conserved. In addition, our functional results reveal contrasting patterns of ERK activation in response to aversive and reward signals, and suggest that activation of the TS is a common feature of psychostimulant drugs.

The use of BAC transgenic mice has been instrumental in improving our understanding of the anatomo-functional organization of the striatum (Bertran-Gonzalez *et al.* 2008; Matamales *et al.* 2009). Recent in-depth analyses of the anatomical arrangement of D2R/A2aR- and D1R-SPNs have revealed that the randomness of the distribution of these two neuronal populations varied along the rostro-caudal axis (Gangarossa *et al.* 2013b; Gagnon *et al.* 2017; Ren *et al.* 2017). In fact, while D2R/A2aR- and D1R-SPNs are randomly distributed in the majority of the dorsal striatum (Gangarossa *et al.* 2013b; Ren *et al.* 2017; Wei *et al.* 2018), a segregated anatomical organization has been unveiled in the TS (Gangarossa *et al.* 2013b). We analyzed the distribution of eGFP and RFP-positive cells in the TS in combination with specific markers of D2R- and D1R-SPNs, and extending our previous observations (Gangarossa *et al.* 2013b), we confirmed the existence of a D2R/A2aR-lacking area composed exclusively of D1R-expressing SPNs. In addition, the use of *Drd2*-eGFP/*Drd1a*-tdTomato mice allowed us to delineate two additional TS territories. The first one, juxtaposed to the D2R/A2aR-lacking area, corresponds to a large territory in which D1R- and D2R-SPNs are evenly intermingled. The direct visualization of D1R- and D2R-SPNs revealed that in the intermingled area, SPNs co-expressing both D1R and D2R were preferentially clustered adjacently to the corpus callosum. This contrasts with the homogenous distribution previously described in the anterior dorsal striatum (Gagnon *et al.* 2017; Ren *et al.* 2017; Wei *et al.* 2018). The second territory, embedded between the striatum and the central amygdala is characterized by a high percentage of D1R/D2R-SPNs (∼33%). This region, which most likely corresponds to the amygdalostriatal transition (AST) area (Fudge and Haber 2002; Jolkkonen *et al.* 2001), is the striatal territory comprising the highest density of D1R/D2R-SPNs, together with the bundle-shaped area of the caudomedial shell of the nucleus accumbens (Gangarossa *et al.* 2013a; Gagnon *et al.* 2017). Although the connectivity and function of this striatal population remain to be established, recent data indicate that D1R/D2R-SPNs display unique morphological features (Gagnon *et al.* 2017) and molecular profile (Gokce *et al.* 2016; Saunders *et al.* 2018). Future studies will be required to determine whether TS D1R/D2R-SPNs share similar features or define another SPNs population.

Our comparative analysis demonstrates that D2R/A2aR- and D1R-SPNs distribution is anatomically conserved across the *Muridae* family (*Rodentia* order). Indeed, even though size and shape slightly differed, the D2R/A2aR-lacking area was unambiguously identified in different wild mouse strains, gerbils and rats. Interestingly, recent high-throughput projectome studies revealed that anatomical origins of cortico- and thalamo-striatal sensory inputs innervating the TS are also conversed in both mice and rats (Hintiryan *et al.* 2016; Hunnicutt *et al.* 2016; Jiang and Kim 2018). Indeed, contrasting with the rostral striatum which is mainly innervated by cingulate, motor and somatosensory cortices (Hunnicutt *et al.* 2016), TS SPNs receive convergent excitatory inputs arising exclusively from the visual and auditory cortices and TS-projecting DA neurons from the lateral part the SNc (Menegas *et al.* 2015; Hintiryan *et al.* 2016; Hunnicutt *et al.* 2016; Jiang and Kim 2018; Poulin *et al.* 2018; Deniau *et al.* 1996). Despite anatomical constancy across species, the functional role of the TS has been poorly addressed. Recently two studies revealed that the TS is not essential to promote movement. Indeed, lesion of TS-projecting DA did not affect the time spent moving and the velocity (Menegas *et al.* 2018). Moreover, while direct activation of D1R-SPN in the dorsal striatum (rostral part) using optogenetics promotes motor activity (Kravitz *et al.* 2010), activation of TS D1R-SPNs failed to elicit movement (Guo *et al.* 2018). Instead, D1R-SPNs of the TS appear to play an important role in auditory decisions (Guo *et al.* 2018; Xiong *et al.* 2015) and contribute to reinforcement learning that promotes avoidance of threatening stimuli (Menegas *et al.* 2018). Future studies are warranted to establish whether a causal relationship may exist between the aforementioned functions and the activation of D1R-SPNs within specific TS territories.

Our functional study, assessed by monitoring ERK phosphorylation (i.e. activation), provides important clues concerning the recruitment of SPNs within the TS. While salient and aversive signals are known to activate TS-projecting DA neurons (Menegas *et al.* 2017; Menegas *et al.* 2018), aversive stimuli triggering innate or learned avoidance responses failed to trigger ERK activation in the TS. However, our results suggest that TS SPNs activity might depend on the nature of defensive reactions (freezing vs avoidance/escape) engaged when exposed to an aversive stimulus signaled or not by a predictive cue. Thus, the level of P-ERK in the TS was unaltered in mice displaying freezing behavioral responses, while it decreased in mice that learned to actively avoid an aversive stimulus by initiating a locomotor response. In addition, we have found that the administration of LiCl, an aversive drug that causes nausea and visceral malaise, also blunted ERK activation in the TS. This decreased phosphorylation was particularly evident in the AST and the ventral edge of the intermingled area, which both integrate visceral information (Hintiryan *et al.* 2016). Moreover, ERK phosphorylation remained unchanged following fasting, satiety or food palatability suggesting that the TS appears to be unsensitive to inner state information associated with feeding behavior.

Our survey using various pharmacological agents revealed that TS SPNs are particularly responsive to psychostimulants drugs. Thus, the administration of d-amphetamine, MDMA, cocaine or methylphenidate increased ERK activation selectively in D1R-SPNs of the D2R/A2aR-lacking area and to a lesser extent in the ventral edge of the intermingled area. The similar pattern of ERK phosphorylation induced by these drugs in the TS contrasts with the one observed in the rostral striatum, where d-amphetamine activates ERK and intracellular signaling events preferentially in SPNs located in the striosomal compartment (Graybiel *et al.* 1990; Biever *et al.* 2015). In the TS, a more pronounced activation was observed with d-amphetamine and MDMA compared to cocaine and methylphenidate. The lack of effects of NET and SERT inhibitors (desipramine and fluoxetine, respectively) combined with the increase in ERK phosphorylation following blockade of DA reuptake (GBR12783), which triggered a pattern of ERK activation comparable to cocaine or methylphenidate, strongly suggest that the mechanisms by which extra-synaptic DA is released account for the difference observed between amphetamine and cocaine. Because the TS integrates visual and auditory information, future studies will be required to determine whether activation of TS D1R-SPNs contribute to psychostimulant-induced arousal and alertness.

The activation of ERK in the D1R-SPNs of the TS by d-amphetamine is prevented by SCH23390 indicating that, similarly to the rostral part of the dorsal striatum and the nucleus accumbens, D1R activation is required (Valjent *et al.* 2005; Bertran-Gonzalez *et al.* 2008; Gerfen *et al.* 2008). Interestingly, systemic pharmacological stimulation of D1R, using agonists known to trigger D1R signaling through cAMP production (Liu and Graybiel 1998; Kuroiwa *et al.* 2008), was sufficient to activate ERK in the TS. Given the similarity observed between the patterns of ERK phosphorylation induced by D1R agonists and d-amphetamine, it is tempting to hypothesize that common intracellular mechanisms could account for this activation. Thus, the inhibition of ERK dephosphorylation through the cAMP/PKA/DARPP-32 has been shown to be essential to sustain D1R-mediated ERK phosphorylation in the striatum (Valjent *et al.* 2005; Gerfen *et al.* 2008). The triggering events in D1R-dependent ERK activation, meanwhile, involve synergistic intracellular events. In fact, cAMP-mediated mechanisms involving the PKA-dependent recruitment of Rasgrp2 and/or the activation of the cAMP sensor NCS-Rapgef2 have been recently unveiled (Nagai *et al.* 2016; Jiang *et al.* 2017). Moreover, ERK activation in D1R-SPNs has also been shown to rely on the activation of Fyn which, by phosphorylating GluN2B of NMDAR, leads to an increase of calcium influx and a subsequent activation of the Ras-GRF1/ERK cascade (Fasano *et al.* 2009; Pascoli *et al.* 2011; Jin *et al.* 2019). Such cellular events provide a mechanism through which glutamate contribute to the regulation of ERK activation induced by d-amphetamine and D1R agonists (Choe *et al.* 2002; Valjent *et al.* 2005; Pascoli *et al.* 2011). Because TS-projecting DA neurons express VGLUT2 (Root *et al.* 2016; Poulin *et al.* 2014; Poulin *et al.* 2018), future studies will be necessary to determine whether glutamate co-released by these DA neurons is essential for D1R-dependent stimulation of ERK in the TS.

Interestingly, when intracellular cAMP was increased through PDE10A inhibition, enhanced ERK phosphorylation was restricted to the ventral part of the intermingled area and confined in D2R-SPNs. This is consistent with previous studies showing that PDE10A inhibition increased PKA-dependent phosphorylation levels in D2R-SPNs of the dorsomedial striatum (Polito *et al.* 2015). This difference was demonstrated to result from a powerful action of phosphatase-1 (PP-1), which prevents PKA substrates from remaining phosphorylated, whereas DARPP-32 efficiently inhibits PP-1 in D2R-SPNs (Nishi *et al.* 2008; Polito *et al.* 2015). The DARPP-32/PP-1 loop also regulates ERK phosphorylation (Valjent *et al.* 2005), and we therefore hypothesize that the increased ERK phosphorylation in D2R-SPNs in response to PDE10A inhibition results from the same mechanism.

In conclusion, our study provides anatomical and molecular evidence suggesting that the TS can indeed be considered as a fourth striatal domain. Our functional analysis also indicates that the TS is a major target of psychostimulant drugs. Future investigations are now required to decipher the functional implication of TS in the action of these drugs.

## Acknowledgments

This work was supported by Inserm, Fondation pour la Recherche Médicale (FRM) and The French National Research Agency ANR-DOPAFEAR (EV), ANR-SEXREV (FV). GG is supported by Allen Foundation Inc. (2016.326) and Nutricia Research Foundation (2017-20) grants. The authors thank the animal breeding facility of Montpellier University (CECEMA).

## Supplemental Figure Legends

**Supplemental Figure 1: Distribution of total ERK in the mouse TS.** ERK immunoreactivity (magenta) in the TS of C57Bl/6J mice. Scale bar, 200 µm. TS: tail of the striatum; GPe: external globus pallidus; AST: amygdalostriatal transition; LA: lateral amygdala; CeA: central amygdala.

**Supplemental Figure 2: Regulation of ERK phosphorylation in the TS by aversive stimuli triggering innate avoidance.** (a) P-ERK immunoreactivity (magenta) in the TS of C57Bl/6J mice (intruder) and FVB/N mice (resident). (b) P-ERK immunoreactivity (magenta) in the TS of C57Bl/6J mice exposed to natural stressors (rat bedding or TMT). Scale bar, 200 µm. TS: tail of the striatum; GPe: external globus pallidus; AST: amygdalostriatal transition; LA: lateral amygdala. For P-ERK cell counting see Table 3. For detailed statistics see Supplemental Table 7.

**Supplemental Figure 3: Regulation of ERK phosphorylation in the TS following auditory Pavlovian conditioning.** (a) Time course of the freezing response during the conditioning phase. Mice were exposed to unpredictable mild footshocks (US, 5x) (upper panel, left), auditory stimulus (CS, 5x) (upper panel, right), to paired CS-US (5x) (lower panel, left) or unpaired CS-US (5x) (lower panel, right). Insets in the lower panels indicate the freezing response during the exposure of each CS in the paired (left) and unpaired (right) groups. Data are presented as means ± sem. * p < 0.05, ** p < 0.01, *** p < 0.001 T_13-17min_ vs T_1min_. For detailed statistics see Supplemental Table 11. (b) P-ERK immunoreactivity (magenta) in the TS of C57Bl/6J mice exposed to auditory fear conditioning. Scale bar, 200 µm. TS: tail of the striatum; GPe: external globus pallidus; AST: amygdalostriatal transition; LA: lateral amygdala. For P-ERK cell counting see Table 3. For detailed statistics see Supplemental Table 7 (P-ERK counting) and 10 (behavior).

**Supplemental Figure 4: Food-related behaviors.** (a) Variation of body weight (g, grams) in control (n = 4), fasting (n = 3), refeeding_1cycle_ (n = 4), refeeding_6cycles_ (n = 4) and HFHS_binge_ (n = 4) mice. Note the decrease and the increase in body weight in fasting/refeeding and HFHS groups. Data are presented as means ± sem. *** p < 0.001 all groups vs Control. (b) Food intake (kcal) during the 30 min of food exposure (chow or HFHS diet) before perfusion. Data are presented as means ± sem. *** p < 0.001 all groups vs Control. The fasting group is not plotted in Fig. S4b because this experimental group did not have access to food before perfusion. For detailed statistics see Supplemental Table 11.

**Supplemental Figure 5: Acute d-amphetamine administration increases P-ERK in the TS.** (a) P-ERK immunoreactivity (magenta) in the TS of C57Bl/6J mice 15 min after saline or d-amphetamine (10 mg/kg) administration. Scale bar, 100 µm. (b) Quantification of P-ERK-immunoreactive neurons in the intermingled, the D2R/A2aR-lacking and the AST areas of mice treated with saline (n = 4) or d-amphetamine and perfused 15 (n = 6) or 30 min (n = 3) after injection. Data are presented as means ± sem. ** p < 0.01, *** p < 0.001 sal vs d-amph. GPe: external globus pallidus; AST: amygdalostriatal transition; LA: lateral amygdala; TS: tail of the striatum; sal: saline; d-amph: d-amphetamine. For detailed statistics see Supplemental Table 12.

