## Supplementary material for "Contrasting patterns of ERK activation in the tail of the striatum in response to aversive and rewarding signals": Text

**Running title:** ERK, dopamine and the caudal striatum

**Keywords:** striatum, dopamine, ERK, D1R, D2R, psychostimulants

### Suppl. Table 1 (Statistics of Figure 3)

| Figure 3 | Mice (n) | Slice (n) | Neurons (n) | TS area | Statistical analysis | p-value |
|---|---|---|---|---|---|---|
| Fig. 3b | n=3 | n=5 | SKF38393<br>SKF38393+CGS21680<br>(n=48) | D2R/A2A-lacking | Paired<br>t-test | 0,229 |
| Fig. 3d | n=3 | n=3 | SKF38393<br>SKF38393+CGS21680<br>(n=47) | AST | Paired<br>t-test | <0,001 |

### Suppl. Table 2 (Statistics of Figure 5)

| Figure 5 | Mice (n) | Behavior | Statistical analysis | F-value | p-value |
|---|---|---|---|---|---|
| Fig. 5b | habituation (15)<br>Day 1 (12)<br>Day 2 (6) | habituation | Unpaired<br>t-test | | 0,827 |
| | | CS+ | | | <0,001 |
| | | CS- | | | 0,476 |

| Figure 5 | Mice (n) | Cell-type specificity | TS area | Statistical analysis | F-value | p-value |
|---|---|---|---|---|---|---|
| Fig. 5c | Context (3)<br>Day1 (6)<br>Day2 (6) | None | Intermingled | One-way<br>ANOVA | $F_{(2, 12)} = 5,82$ | <0,05 |
| | | | D2R-lacking | | $F_{(2, 12)} = 8,54$ | <0,01 |
| | | | AST | | $F_{(2, 12)} = 0,55$ | 0,589 |
| Fig. 5d | Saline (4)<br>LiCl (3) | None | Intermingled | Unpaired<br>t-test | | <0,05 |
| | | | D2R-lacking | | | <0,05 |
| | | | AST | | | <0,05 |

### Suppl. Table 3 (Statistics of Figure 6)

| Figure 6 | Mice (n) | Cell-type specificity | TS area | Statistical analysis | F-value | p-value |
|---|---|---|---|---|---|---|
| Fig. 6b | Saline (4)<br>d-amph$_{15min}$ (6)<br>d-amph$_{30min}$ (3) | D2R-negative | Intermingled | One-way ANOVA | $F_{(2, 10)} = 10,38$ | <0,01 |
| | | | D2R-lacking | | $F_{(2, 10)} = 29,66$ | <0,001 |
| | | | AST | | $F_{(2, 10)} = 2,10$ | 0,173 |
| | | D2R-positive | Intermingled | One-way ANOVA | $F_{(2, 10)} = 4,04$ | 0,052 |
| | | | D2R-lacking | | $F_{(2, 10)} = 3,46$ | 0,072 |
| | | | AST | | $F_{(2, 10)} = 10,38$ | <0,01 |
| Fig. 6d | Saline (3)<br>d-amph$_{15min}$ (4) | Nr4a1-negative | Intermingled | Unpaired t-test | | <0,01 |
| | | | D2R-lacking | | | <0,001 |
| | | | AST | | | <0,001 |
| | | Nr4a1-positive | Intermingled | Unpaired t-test | | 0,051 |
| | | | D2R-lacking | | | 0,075 |
| | | | AST | | | 0,721 |
| Fig. 6f | Saline (8)<br>MPH (4)<br>MDMA (4)<br>Cocaine (6) | None | Intermingled | One-way ANOVA | $F_{(3, 18)} = 34,31$ | <0,001 |
| | | | D2R-lacking | | $F_{(3, 18)} = 19,09$ | <0,001 |
| | | | AST | | $F_{(3, 18)} = 3,03$ | 0,056 |
| Fig. 6g | Saline (8)<br>MPH (4)<br>MDMA (4)<br>Cocaine (6) | D2R-negative | Intermingled | One-way ANOVA | $F_{(3, 18)} = 31,97$ | <0,001 |
| | | | D2R-lacking | | $F_{(3, 18)} = 18,82$ | <0,001 |
| | | | AST | | $F_{(3, 18)} = 7,92$ | <0,01 |
| | | D2R-positive | Intermingled | One-way ANOVA | $F_{(3, 18)} = 1,17$ | 0,349 |
| | | | D2R-lacking | | $F_{(3, 18)} = 0,99$ | 0,418 |
| | | | AST | | $F_{(3, 18)} = 0,93$ | 0,445 |

### Suppl. Table 4 (Statistics of Figure 7)

| Figure 7 | Mice (n) | Cell specificity | TS area | Statistical analysis | F-value | p-value |
|---|---|---|---|---|---|---|
| Fig. 7b | Saline (3)<br>GBR (3)<br>Desipramine (4)<br>Fluoxetine (4) | None | Intermingled | One-way ANOVA | $F_{(3, 10)} = 9,51$ | <0,01 |
| | | | D2R-lacking | | $F_{(3, 10)} = 43,65$ | <0,001 |
| | | | AST | | $F_{(3, 10)} = 2,80$ | 0,095 |

### Suppl. Table 5 (Statistics of Figure 8)

| Figure 8 | Mice (n) | Cell specificity | TS area | Statistical analysis | F-value | p-value |
|---|---|---|---|---|---|---|
| Fig. 8b | d-amph (6)<br>d-amph+SCH (4) | None | Intermingled | Unpaired t-test | | <0,001 |
| | | | D2R-lacking | | | <0,001 |
| | | | AST | | | <0,05 |
| Fig. 8d | Sal (3)<br>SKF81297 (3)<br>SKF83822 (4)<br>SKF82595 (3) | None | Intermingled | One-way ANOVA | $F_{(3, 9)} = 15,81$ | <0,001 |
| | | | D2R-lacking | | $F_{(3, 9)} = 13,45$ | <0,001 |
| | | | AST | | $F_{(3, 9)} = 6,06$ | <0,05 |

### Suppl. Table 6 (Statistics of Figure 9)

| Figure 9 | Mice (n) | Cell specificity | TS area | Statistical analysis | F-value | p-value |
|---|---|---|---|---|---|---|
| Fig. 9b | Saline (4) Papaverine (3) | None | Intermingled | Unpaired t-test | | <0,05 |
| | | | D2R-lacking | | | 0,360 |
| | | | AST | | | 0,420 |
| Fig. 9c | Saline (4) Papaverine (3) | D2R-negative | Intermingled | Unpaired t-test | | 0,909 |
| | | | D2R-lacking | | | 0,286 |
| | | | AST | | | 0,210 |
| | | D2R-positive | Intermingled | Unpaired t-test | | <0,05 |
| | | | D2R-lacking | | | 0,846 |
| | | | AST | | | 0,642 |
| Fig. 9e | Saline (4) Rolipram (4) | D2R-negative | Intermingled | Unpaired t-test | | 0,059 |
| | | | D2R-lacking | | | 0,269 |
| | | | AST | | | <0,01 |
| | | D2R-positive | Intermingled | Unpaired t-test | | 0,223 |
| | | | D2R-lacking | | | 0,097 |
| | | | AST | | | 0,639 |

### Suppl. Table 7 (Statistics of Table 3)

| Table 3 | Mice (n) | Cell specificity | TS area | Statistical analysis | F-value | p-value |
|---|---|---|---|---|---|---|
| 3.1 | Intruder (3) Resident (3) | None | Total | Unpaired t-test | | 0,389 |
| | | | Intermingled | | | 0,447 |
| | | | D2R-lacking | | | 0,219 |
| | | | AST | | | 0,053 |
| 3.2 | Mouse (3) Rat (3) | None | Total | Unpaired t-test | | 0,832 |
| | | | Intermingled | | | 0,687 |
| | | | D2R-lacking | | | 0,422 |
| | | | AST | | | 0,069 |
| 3.2 | Water (4) TMT (3) | None | Total | Unpaired t-test | | 0,293 |
| | | | Intermingled | | | 0,737 |
| | | | D2R-lacking | | | 0,315 |
| | | | AST | | | 0,087 |
| 3.3 | Context (3) CS (6) US (6) CS-US paired (6) CS-US unpaired (3) | None | Total | One-way ANOVA | $F_{(4, 19)} = 1,63$ | 0,207 |
| | | | Intermingled | | $F_{(4, 19)} = 1,72$ | 0,187 |
| | | | D2R-lacking | | $F_{(4, 19)} = 2,68$ | 0,063 |
| | | | AST | | $F_{(4, 19)} = 6,18$ | 0,666 |

### Suppl. Table 8 (Statistics of Table 4)

| Table 4 | Mice (n) | Cell specificity | TS area | Statistical analysis | F-value | p-value |
|---|---|---|---|---|---|---|
| 4.1 | Control (4)<br>Fasting (3)<br>$Refeeding_{1cycle}$ (4)<br>$Refeeding_{6cycles}$ (4)<br>$HFHS_{binge}$ (4) | None | Total | One-way ANOVA | $F_{(4, 14)}$ = 2,68 | 0,075 |
| | | | Intermingled | | $F_{(4, 14)}$ = 2,13 | 0,130 |
| | | | D2R-lacking | | $F_{(4, 14)}$ = 1,49 | 0,257 |
| | | | AST | | $F_{(4, 14)}$ = 2,40 | 0,099 |

### Suppl. Table 9 (Statistics of Table 5)

| Table 5 | Mice (n) | Cell specificity | TS area | Statistical analysis | F-value | p-value |
|---|---|---|---|---|---|---|
| 5.1 | Control (7)<br>DOI (4)<br>PCP (3)<br>MK801 (3) | None | Total | One-way ANOVA | $F_{(3, 13)}$ = 10,61 | <0,001 |
| | | | Intermingled | | $F_{(3, 13)}$ = 7,28 | <0,01 |
| | | | D2R-lacking | | $F_{(3, 13)}$ = 5,83 | <0,01 |
| | | | AST | | $F_{(3, 13)}$ = 3,44 | <0,05 |
| 5.2 | Saline (3)<br>PTZ (3)<br>Kainate (3) | None | Total | One-way ANOVA | $F_{(2, 6)}$ = 4,69 | 0,057 |
| | | | Intermingled | | $F_{(2, 6)}$ = 0,57 | 0,595 |
| | | | D2R-lacking | | $F_{(2, 6)}$ = 1,75 | 0,252 |
| | | | AST | | $F_{(2, 6)}$ = 11,51 | <0,01 |

### Suppl. Table 10 (Statistics of Suppl. Figure S3)

| Figure S3 | Mice (n) | Behavior | Statistical analysis | F-value | p-value |
|---|---|---|---|---|---|
| S3a | Shock US (6)<br>Sound CS (6)<br>CS-US paired (6)<br>CS-US unpaired (3) | US | One-way ANOVA | $F_{(13, 70)}$ = 33,85 | <0,001 |
| | | CS | | $F_{(13, 70)}$ = 0,78 | 0,674 |
| | | CS-US paired | | $F_{(13, 70)}$ = 29,30 | <0,001 |
| | | CS-US unpaired | | $F_{(16, 34)}$ = 9,83 | <0,001 |

### Suppl. Table 11 (Statistics of Suppl. Figure S4)

| Figure S4 | Mice (n) | Behavior | Statistical analysis | F-value | p-value |
|---|---|---|---|---|---|
| S4a | Control (4)<br>Fasting (3)<br>$Refeeding_{1cycle}$ (4)<br>$Refeeding_{6cycles}$ (4)<br>$HFHS_{binge}$ (4) | Variation of body weight | One-way ANOVA | $F_{(4, 14)}$ = 87,90 | <0,001 |

| | | | | | |
|---|---|---|---|---|---|
| S4b | Control (4)<br>$Refeeding_{1cycle}$ (4)<br>$Refeeding_{6cycles}$ (4)<br>$HFHS_{binge}$ (4) | Food intake | One-way ANOVA | $F_{(3, 12)} = 102{,}40$ | <0,001 |

### Suppl. Table 12 (Statistics of Suppl. Figure S5)

| Figure S5 | Mice (n) | Cell-type specificity | TS area | Statistical analysis | F-value | p-value |
|---|---|---|---|---|---|---|
| S5b | Saline (4)<br>d-amph$_{15min}$ (6)<br>d-amph$_{30min}$ (3) | None | Intermingled | One-way ANOVA | $F_{(2, 10)}$ = 10,80 | <0,01 |
| | | | D2R-lacking | | $F_{(2, 10)}$ = 29,17 | <0,001 |
| | | | AST | | $F_{(2, 10)}$ = 1,43 | 0,283 |

**Supplemental Figure Legends**

**Supplemental Figure 1: Distribution of total ERK in the mouse TS.** ERK immunoreactivity (magenta) in the TS of C57Bl/6J mice. Scale bar, 200 µm. TS: tail of the striatum; GPe: external globus pallidus; AST: amygdalostriatal transition; LA: lateral amygdala; CeA: central amygdala.
