## Supplementary figures and images for "Contrasting patterns of ERK activation in the tail of the striatum in response to aversive and rewarding signals"

### Suppl. Figure 1

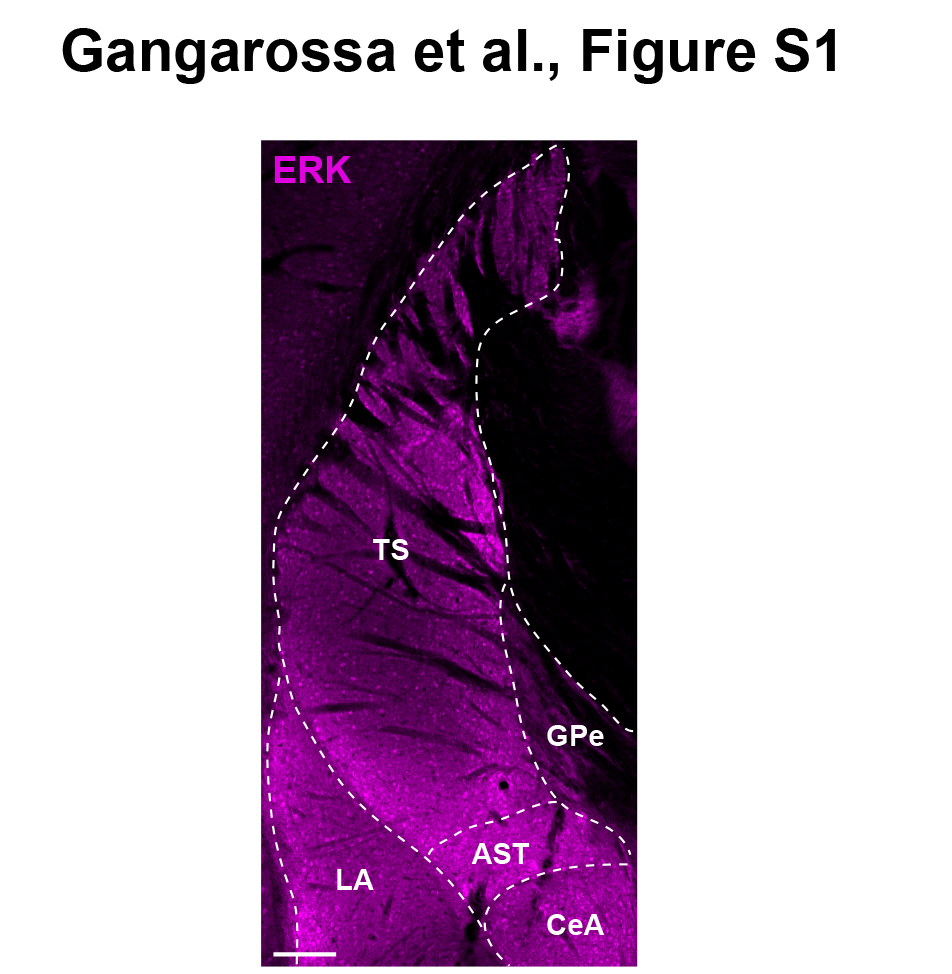

### Suppl. Figure 2

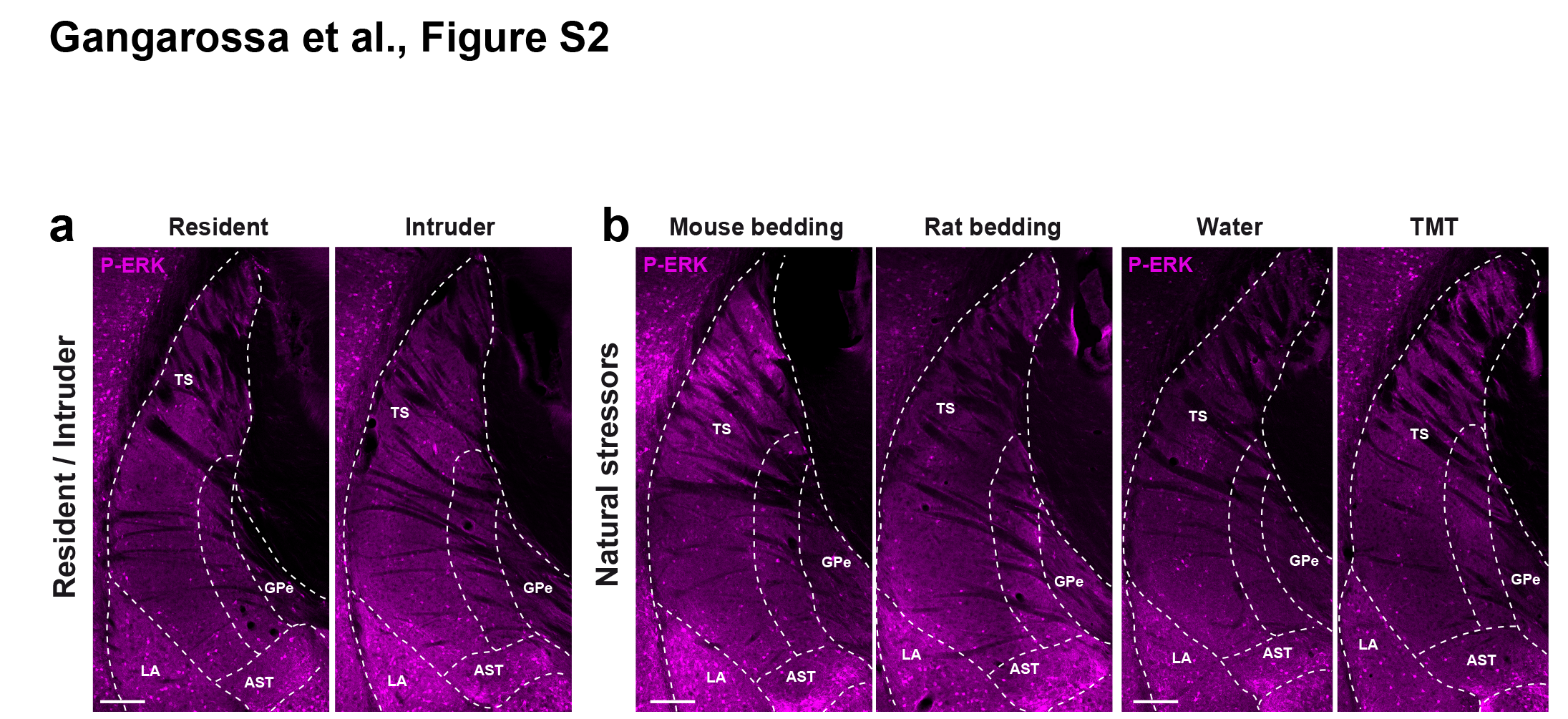

### Suppl. Figure 3

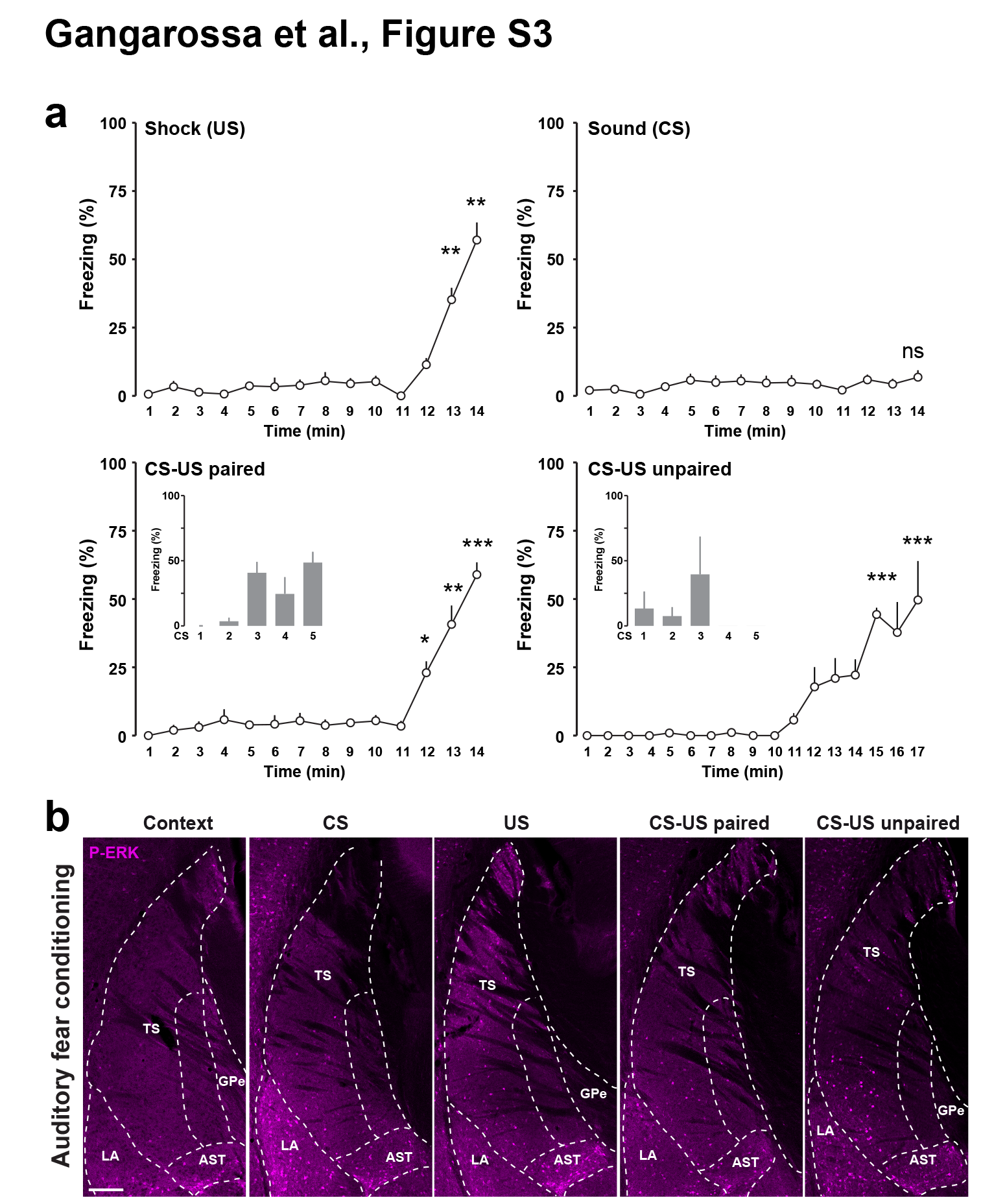

### Suppl. Figure 4

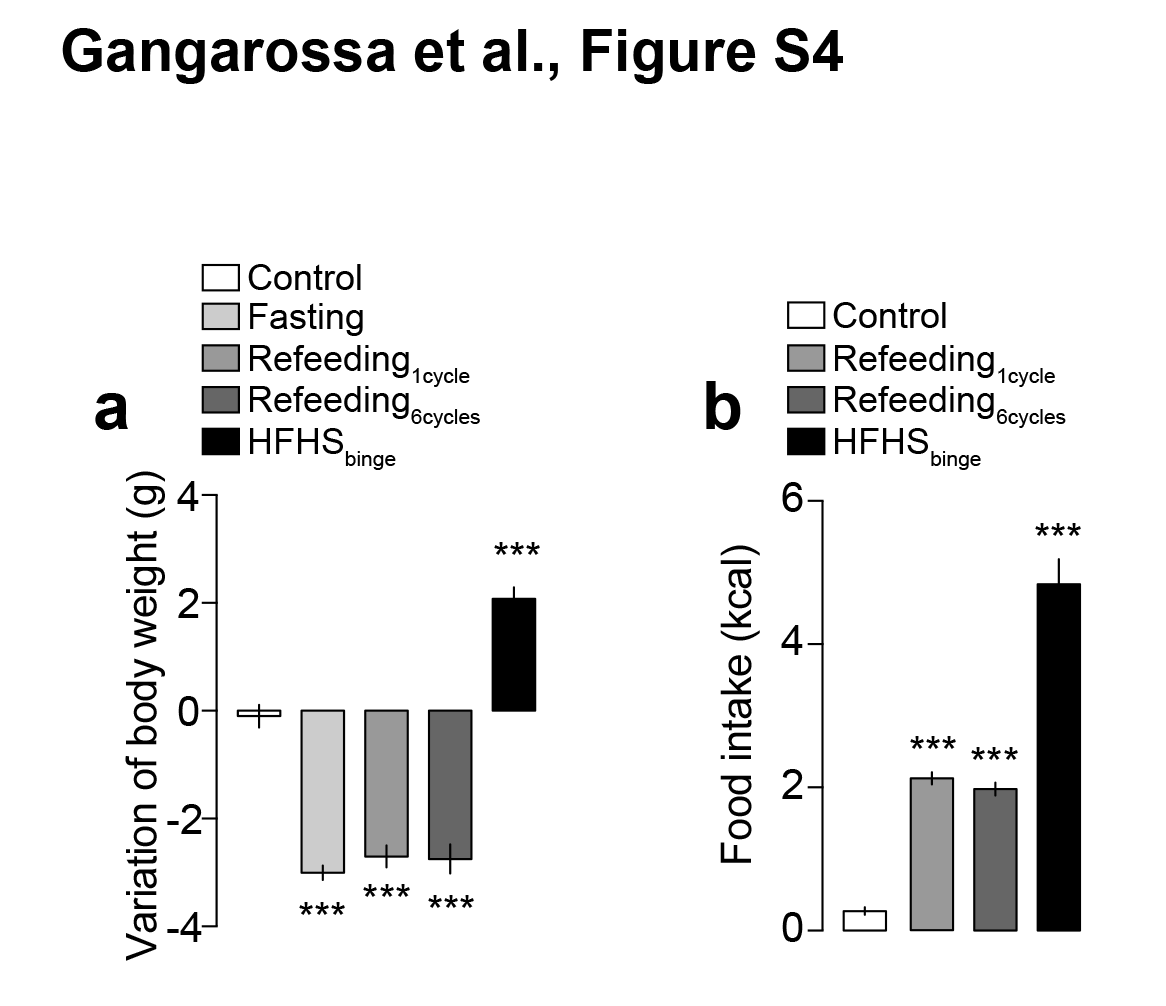

### Suppl. Figure 5

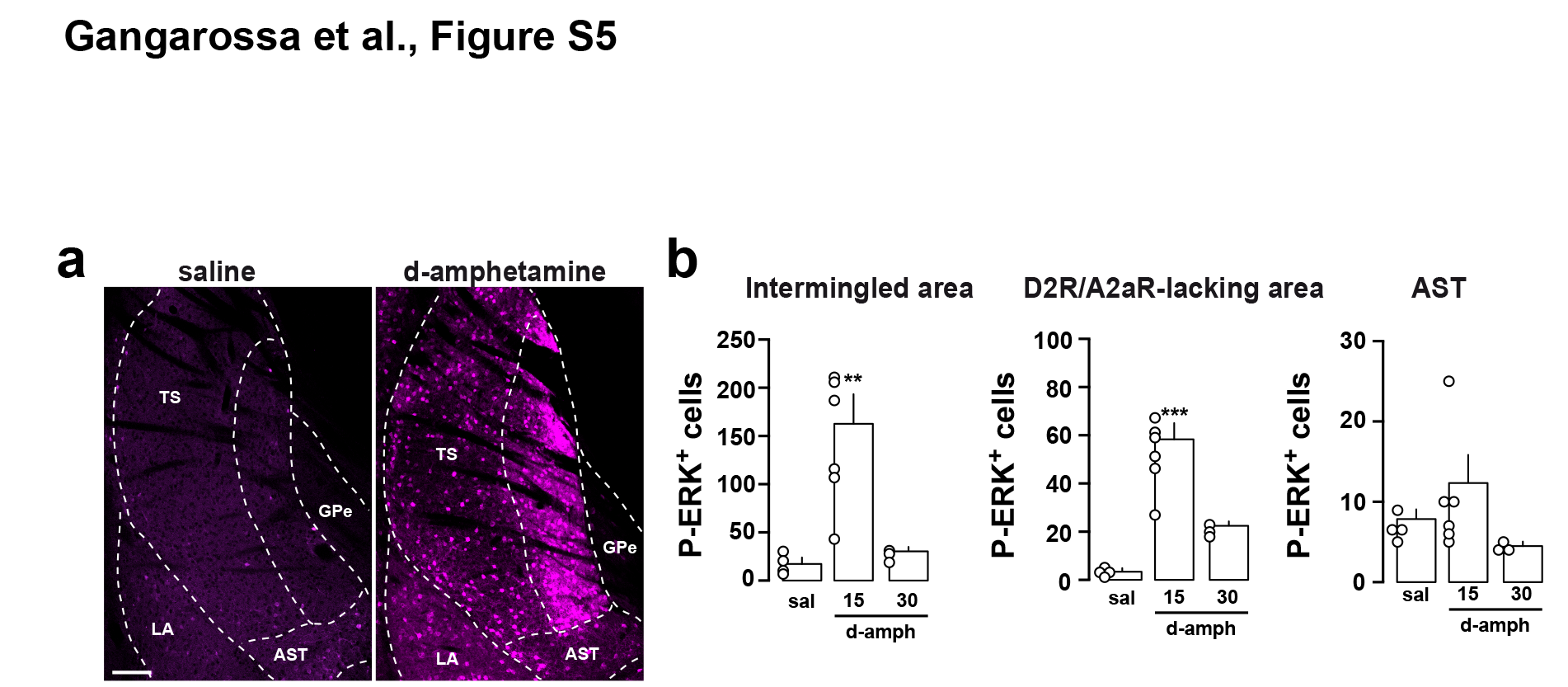
